# IP-MS Analysis of ESX-5 and ESX-1 Substrates Enables Mycobacterial Species Identification

**DOI:** 10.1101/2020.06.07.138784

**Authors:** Qingbo Shu, Meena Rajagopal, Jia Fan, Lingpeng Zhan, Xiangxing Kong, Yifan He, Suwatchareeporn Rotcheewaphan, Christopher J. Lyon, Wei Sha, Adrian M. Zelazny, Tony Hu

**Author notes:** **Correspondence:** Tony Hu.

## Abstract

Pulmonary disease resulting from non-tuberculous mycobacteria (NTM) infection has emerged as an increasingly prevalent clinical entity in the past two to three decades, but there are no standardized, commercial assays available for the molecular diagnosis of NTM infections from clinical samples. Herein we discuss the development of an assay that employs immunoprecipitation coupled with mass spectrometry (IP-MS) to rapidly and accurately discriminate prevalent slow-growing mycobacterial species (i.e., *M. avium* and *M. intracellulare, M. kansasii, M. gordonae, M. marinum* and *M. tuberculosis*) during early growth in mycobacterial growth indicator tube (MGIT) cultures. This IP-MS assay employs antibodies specific for conserved tryptic peptides of M. tuberculosis EsxN (AQAASLEAEHQAIVR) and CFP-10 (TQIDQVESTAGSLQGQWR) to capture and identify specific mycobacterial species and allows species-specific mycobacterium identification at the first sign of MGIT culture positivity.

## Introduction

The genus *Mycobacterium* contains more than 190 known species (1), including *M. tuberculosis* (*Mtb*), the leading cause of death from infectious disease, and is primarily spread by respiratory droplet exposure through contact with infected individuals. NTM, however, are often encountered during exposure to environmental sources and may cause a variety of disease manifestations, including respiratory, lymph node, skin, soft tissue and disseminated infections. Some NTM species are primarily associated with pulmonary disease, including *M. abscessus, M. kansasii*, and species within the *M. avium* complex (MAC), which includes *M. avium* and *M. intracellulare*. Distribution of these NTM species show geographic difference, and MAC, *M. gordonae, M. xenopi*, and *M. kansasii* and rapid growers including *M. abscessus* and *M. fortuitum* are the most prevalent NTM species worldwide (2). *M. gordonae* and *M. xenopi* isolations were limited to distinct geographical regions; mainly in Europe and Ontario. The most common mycobacterial organism associated with pulmonary disease in the United States is *Mycobacterium avium* complex (MAC), followed by *Mycobacterium kansasii* (both slowly growing mycobacteria) and *Mycobacterium abscessus* (rapidly growing species) (3, 4).

Due to the difficulty of obtaining diagnostically useful specimens from some patients, serodiagnostic assays that detect antibodies against specific mycobacterial antigens are promising alternatives, and such assays have been investigated for their ability to detect MAC and *M. abscessus* antigens. However, the potential for such antibodies to cross-react with homologous proteins produced by different mycobacterial species can result in inconclusive or erroneous identification of the mycobacterial species responsible for a suspected infection (5).

Tuberculin skin tests and interferon-gamma release assays (IGRAs) that assess *Mtb* exposure based on the patient’s immune response upon exposure to Mtb factors are widely used to diagnose *Mtb* infection. However, similar assays do not exist for common NTM species.

IGRAs are employed to diagnose individuals with suspected *Mtb* infection by quantifying the immune responses samples spiked with the abundantly secreted ESX-1 substrates ESAT-6 (EsxA) and CFP-10 (EsxB). These antigens are primarily secreted by highly related Mtb complex species, but may also be secreted by some NTM species that do not belong to this complex, including *M. kansasii* and *M. marinum*. However, most mycobacteria secrete Antigen 85 complex proteins via the TAT system and secrete EsxN via the ESX-5 system, which may allow these proteins to serve as potential biomarkers for more broad-spectrum diagnosis of NTM infections.

Culture remains the gold standard for laboratory confirmation of NTM infection, species identification, and drug susceptibility testing (6), and various automated culture-reading systems (e.g. BD Bactec MGIT) are employed for rapid evaluation of mycobacterial growth in liquid culture. NTM species identification is typically performed using approaches that employ molecular probes or PCR gene sequencing (7) or MALDI-TOF MS analysis (8, 9), although subculture required for MALDI-TOF MS analysis takes several days to weeks to obtain sufficient biomass for species identification, and the efficiency of the protein extraction procedure can also affects identification accuracy. The major drawback of *secA1* gene sequencing for species identification is that it cannot discriminate members of the MTB complex, which share the same *secA1* gene sequence. With the increasing knowledge of *Mtb* sub-linages and strains, it becomes useful to identify distinct *Mtb* sub-linages. Also, it will be useful to distinguish *M. bovis* BCG with *Mtb* using molecular assays.

Moreover, the isolation of organisms of the *Mtb* complex using solid culture plate needs biosafety level 3 (BSL-3) facilities, which dramatically limits the application of this method in a common clinical lab. Therefore, this bacterial colony-based identification method needs to address issues with sample preparation though it has been widely applied. The bacterial pellet-based method avoids time-consuming subculture; however, it still requires BSL-3 facilities to complete sample preparation. Most importantly, it suffers from low success rates specially with slowly growing mycobacteria, in part due to insufficient amount of bacterial biomass in the sample. Another challenge it faces with is low rate of identification when MGIT tubes were polymicrobial, where isolation of *Mtb* was mixed with NTM or other bacteria on subculture. We therefore proposed to use the MGIT filtrates and detect two bacterial antigen derived peptides by parallel-reaction monitoring (PRM) in order to avoid time-consuming subculture, processing of samples in BSL-3 facilities and address the low identification rate in polymicrobial samples. Liquid chromatography coupled tandem mass spectrometry (LC-MS/MS) in PRM mode (PRM-MS) has been used to discriminate slow-growing mycobacterial species by identifying the species of origin of abundantly secreted mycobacterial proteins (e.g. Ag85B) in mycobacterial liquid culture filtrate protein (CFP) samples (10). However, MGIT CFP samples contain abundant protein additives that may interfere with the ability to detect mycobacterial proteins (11). Immunoprecipitation (IP) can be used to enrich target mycobacterial peptides from those produced by abundant proteins that could mask their signal. In order to distinguish highly related proteins from different species, however, it is advisable to use a capture antibody that binds a conserved region of a target peptide and employ mass spectrometry to identify amino acid differences that can distinguish its species of origin.

In this study, we employed LC-MS/MS to analyze secreted proteins present in CFP samples of most common mycobacterial species causing human respiratory infections. *M. marium* is the most common atypical *Mycobacterium* that causes human infections (12). We included *M. marium* as a targeted species as well. This analysis found that tryptic peptides of the ESX-5 substrate EsxN and the ESX-1 substrate CFP-10 were detectable in tryptic digests of CFP samples isolated from liquid cultures, and that tryptic peptides derived from these two proteins and identified via PRM-MS could distinguish all the analyzed mycobacterial species.

## Experimental Procedures

### Experimental Design and Statistical Rationale

Single CFP samples were isolated from MGIT cultures of *M. avium* and *M. intracellulare, M. kansasii* and *M. tuberculosis* and analyzed by LC-MS/MS without replication to identify proteins that were abundantly secreted by each species, after ultracentrifugation with 50kDa filter to remove abundant high molecular weight culture additives and isolate mycobacterial proteins < 50 kDa for LC-MS/MS analysis. The samples are relatively simple in their protein composition, thus one LC-MS/MS run with 2hrs gradient will achieve identification of most proteins. To cover the major population infected with mycobacteria, this study focuses on the most prevalent mycobacterial species, though the proposed method can also target other species with less prevalence. Therefore, the proportions of different mycobacterial species among the 74 patients reflect their prevalence in US. Five patients have biological duplicates of the MGIT samples, and one patient has biological triplicates of the MGIT samples. These replicates serve to validate the method reproducibility. In the US, the rate of Mtb infection in 2010 was 3.6 cases/100,000 persons (13), while the average annual prevalence of pulmonary nontuberculous mycobacteria from 1997 to 2007 was 31 cases/100,000 persons and varied with geography (14). Given the US population as 309.3 million, there are an estimated 11,135 individuals with *Mtb* infections and 95,883 with NTM infections. A study with a 95% confidence level and a confidence interval of 4, would therefore require estimated sample sizes of 570 *Mtb* cases and 593 NTM cases [https://www.surveysystem.com/sscalc.htm]. Geographical variance of prevalence of NTM infection makes it difficult to collect such a number of MGIT specimens from single hospital to accurately represent the prevalence of *Mtb*/NTM infection. However, the major aim of this study was to discover peptide targets that demonstrate the ability to discriminate between mycobacterial species and the potential for translation to clinical settings. Therefore, 74 mycobacterial infection patients are sufficient for method validation though they cannot represent the US population with high statistical power. Ongoing study will increase the number of patients with more geographical variance to validate this method.

### BLAST analysis of Mtb CFP10 and EsxN protein sequence

Reviewed UniProt KB database entries for full-length Mtb CFP10 (P9WNK5) and EsxN (P9WNJ3) protein sequence were searched against the UniProt KB bacterial proteome database using the UniProt BLAST function to identify homologous proteins, using default search parameters (E-Threshold of 10, auto matrix selection, no filtering for low-complexity regions, and permitting gaps in the sequence alignments).

### Liquid culture of mycobacteria

Culture stocks for *M. abscessus* ATCC 19977, *M. avium* ATCC 25291, *M. intracellulare* ATCC 13950 and *M. kansasii* ATCC 12478 were purchased from ATCC, inoculated into 10 mL Middlebrook 7H9 broth supplemented with 10% ADC, 0.2% glycerol, and 0.05% Tween 80, and grown to saturation with 100 rpm rotation in a 37°C incubator. Media was collected at different times post-inoculation for *M. abscessus* (2 days), *M. avium* (21 days), *M. intracellulare* (10 days), and *M. kansasii* (7 days) according to their differential growth rates. Saturated mycobacterial cultures were pelleted by centrifugation at 3,000 ×g for 5 minutes, and supernatants were transferred into a sterile syringe and filtered through a 0.2 µm PES syringe filter and resulting CFP samples were stored at 4°C until use. *M. tuberculosis* CFP was obtained from BEI Resources (NR-14826). *M. abscessus, M. avium, M. intracellulare*, and *M. kansasii* CFP were provided by Dr. Shelley Haydel at the Biodesign Institute, ASU.

### Protein filtration and digestion

Stored NTM samples were passed through a 50 kDa filter unit (Amicon, Thermo) by 14,000 ×g centrifugation at 4 °C, analyzed for protein concentration by BCA assay (Thermo Pierce). For NTM samples, 100 µg of this filtrate was reduced with 10 mM dithiothreitol (DTT, Sigma, USA) at 37 °C for one hour, alkylated with 50 mM iodoacetamide (Sigma, USA) at 25°C for one hour, and then digested with sequencing grade trypsin (Promega, USA) at a protein:enzyme ratio of 50:1 for 16 hours at 37°C. *Mtb* CFP obtained from BEI Resources protein additives was pre-concentrated (15), but otherwise processed in the same manner as all other CFP samples.

### LC-MS/MS analysis of CFP

Trypsin generated peptide mixtures were separated with a linear gradient of 2-37% buffer B (100% ACN and 0.1% formic acid) at a flow rate of 300 nL/min on an EASY-Spray™ C18 LC column (15 cm × 75 µm I.D., 3 µm particle size, Thermo Scientific). An UltiMate 3000 nanoLC system (Thermo Scientific) was on-line coupled to an Orbitrap Velos Pro (NTM samples) or Orbitrap Fusion Lumos instrument (Mtb sample) (Thermo Scientific). MS data were acquired in a data-dependent strategy selecting the fragmentation events based on the precursor abundance in the survey scan (275–1850 Th for NTM samples and 375–1700 Th for the Mtb sample). The resolution of the survey scan was 120,000 at m/z 400 Th.

For the NTM samples, low-resolution CID MS/MS spectra were acquired in rapid CID scan mode for the 15 most intense peaks from the survey scan. A normalized collision energy of 35 was used for fragmentation. Dynamic exclusion was 40 s. The isolation window for MS/MS fragmentation was set to 2 Th. Resulting CID MS/MS spectra were searched for mycobacterial proteomes using Proteome Discoverer (version 2.4, Thermo Scientific) with the following parameters: tryptic digestion, with no more than 2 missed cleavage sites, precursor and product mass tolerances of 10 ppm and 0.6 Da respectively, cysteine carbamidomethylation as stable modification; and methionine oxidation as a dynamic modification. Uniprot Proteome ID for each NTM species used in database searching was, UP000017786 for M. kansasii strain ATCC12478, UP000007137 for *M. abscessus* strain ATCC19977, UP000008004 for *M. intracellulare* strain ATCC 13950, UP000019908 for *M. avium subsp. avium* 2285 (R) and UP000001584 for *Mtb* strain H37Rv. All of them are released on January 15, 2020.

For the *Mtb* sample, HCD MS/MS spectra were acquired in rapid ion trap scan rate and data dependent mode was set as cycle time. A normalized collision energy of 30 was used for fragmentation. Dynamic exclusion was 60 s. The isolation window for MS/MS fragmentation was set to 1.6 Th. The HCD MS/MS spectra were searched against the TB proteome in UniProt using Proteome Discoverer (version 2.4, Thermo Scientific) with the following parameters: tryptic digestion, allowing no more than 2 missed cleavage sites, precursor and product mass tolerances of 10 ppm and 0.02 Da respectively, cysteine carbamidomethylation as a stable modification; methionine oxidation as a dynamic modification.

### IP-MS analysis of culture media and MGIT samples

#### MGIT samples

This research project was reviewed and approved by the Institutional Review Board (IRB) at Tulane University prior to study initiation. Mycobacterial cultures were obtained from patients enrolled in IRB-approved protocols at the National Institutes of Health Clinical Center. Respiratory specimens were decontaminated using the N-acetyl-L-cysteine/sodium hydroxide method, concentrated by 15 min centrifugation at 3,000 × g, and pellets were suspended in 0.8 ml of sterile phosphate buffer saline (PBS). Microscopic examination for the presence of acid-fast bacilli (AFB) was performed using auramine-rhodamine staining (Becton Dickinson, Sparks, MD). All patient samples were inoculated into Bactec 960 MGIT tubes containing the manufacturer’s growth supplement (PANTA) and Middlebrook 7H11 agar (Remel, Lenexa, KS). MGIT samples that were positive for growth were stained with auramine-rhodamine and gram stain to detect the presence of AFB and/or contaminants and sub-cultured on solid media for species identification, and 3 ml of positive MGIT culture was filtered sterilized by passage through a 0.22 µm filter and stored at −80 °C for subsequent LC-MS/MS analysis. Mycobacteria isolated from MGIT cultures were either directly identified by secA gene sequencing (8) and/or after subculture on solid media using matrix-assisted laser desorption/ionization time-of-flight mass spectrometry (MALDI-TOF MS) analysis (9). Species level differentiation within the Mtb complex was determined by a multiplex PCR (10).

#### MGIT and culture filtrate sample digestion

500 µL of MGIT media from patients’ isolates and 500 µL of CFP solutions from M. abscessus ATCC 19977, M. avium ATCC 25291, M. intracellulare ATCC 13950 and M. kansasii ATCC 12478 culture were digested with 5 µg of sequencing grade trypsin after adding 10 µL of 1 M Tris solution. Samples were incubated at 37°C for 16 hours with tilting and rotation on Hulamixer (Thermo Fisher Scientific). After digestion, 5 µL of 10% trifluoracetic acid solution was added into each sample to adjust the pH to 7.

#### Stable isotope-labeled internal standards (IS)

Six peptides (Table 1) labeled with stable isotope arginine (13C615N4) were purchased from New England Peptides (Gardner, MA), and dissolved with 2% acetonitrile, 0.1% trifluoracetic acid at a concentration of 50 µg/mL each to generate an IS peptide mixture. Prior to target peptide immunoprecipitation with antibody conjugated beads, trypsin digested MGIT CFP samples were spiked with 40 µL of diluted IS solution (50 ng/mL) to permit accurate quantification of the isolated target peptides.

**Table 1.**
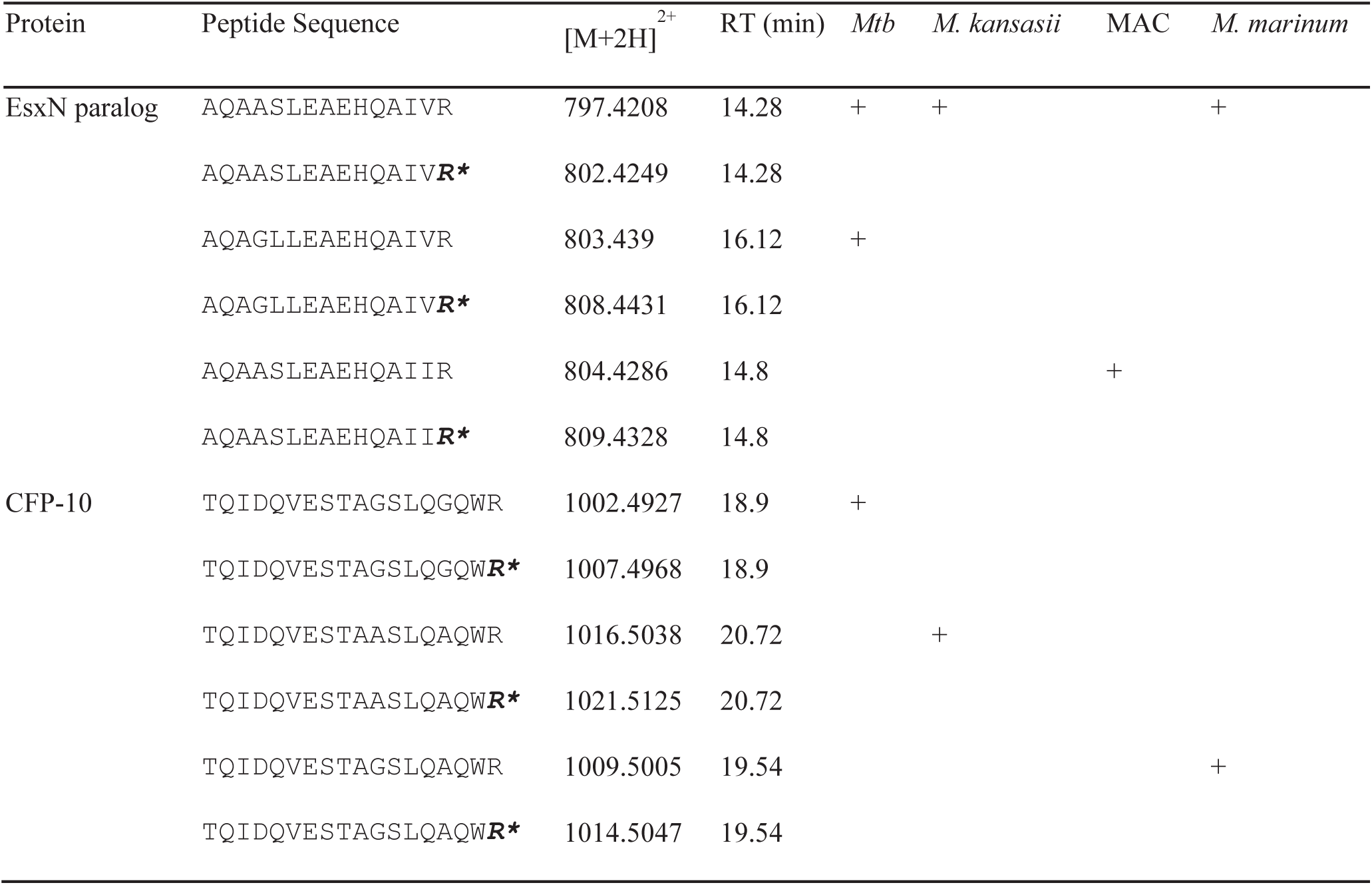
Peptide targets used in PRM-MS analysis for mycobacterial species identification.

#### *Mtb* CFP10 and EsxN peptide antibodies and beads

Custom rabbit polyclonal antibodies (GenScript, Nanjing, China) raised against tryptic peptides of *Mtb* EsxN (AQAASLEAEHQAIVR) and Mtb CFP-10 (TQIDQVESTAGSLQGQWR) were suspended in PBS buffer (pH 7.4) separately, and 40 µg of antibody was bound to 3 mg of protein G Dynabeads (Thermo Fisher Scientific) in 400 µL of PBS buffer containing 0.2% (v/v) Tween-20 binding buffer (0.05 M Tris pH 7.4, 0.15 M sodium chloride, 0.2% n-Dodecyl β-D-maltoside; all reagents from Sigma), which were then washed twice with 200 µL of PBST binding buffer, and suspended in 50 µL of binding buffer.

#### EsxN and CFP-10 peptide immunoprecipitation

EsxN and CFP-10 peptide antibody beads (150 µg each) were incubated with trypsin-digested CFP samples (X µg each) for 1 hour at 25°C on a Hulamixer (Thermo Fisher Scientific) sequentially, then washed three times with 50 µL of binding buffer, and two times with 50 µL of LC grade water (Fisher Scientific). Beads were then incubation for 30 min at 25°C with 100 µL of a 1% (v/v) formic acid solution and precipitated by magnetic isolation, after which supernatants containing eluted peptides loaded onto StageTips (9) generated by packing 200 µL pipette tips with four layers of Empore C8 solid phase extraction disk (3M; Cat. # 2214-C8). StageTips were washed by 25°C centrifugation at 1000g for 3 min with 50 µL of 0.1% (v/v) TFA acetonitrile solution and 50 µL of 0.1% (v/v) TFA prior to sample loading. Each peptide sample was split and loaded onto two StageTips, and captured peptides were eluted with 50 µL of 0.1% (v/v) TFA acetonitrile solution. Paired eluents were combined, dried by vacuum concentration, re-dissolved with 8 µL of sampling buffer containing 0.1% (v/v) formic acid, 2% (v/v) acetonitrile, and centrifuged at 25°C and 21,000 g, for 10 min.

#### PRM-MS analysis

PRM-MS analysis of captured peptides was performed using an QExactive HF-X Orbitrap mass spectrometer coupled with an UltiMate 3000 ultrahigh-pressure liquid chromatography (UHPLC) system (Thermo Scientific). Peptides were loaded onto an Acclaim PepMap100 C18 trap column (300 µm ID × 5 mm, 5 µm, Thermo Fisher Scientific; Cat. # 160454) and then separated on a PepMap RSLC C18 analytical column (75 µm ID×15 cm, 3 µm, Thermo Fisher Scientific, Cat. # 164568). Peptides were eluted with a 300 nL/min gradient generated by mixing buffer A (0.1% formic acid in water) and buffer B (0.1% formic acid in acetonitrile) as follows: a 5 min wash with 5% buffer B, a 17 min 5-38% buffer B gradient, a 2 min 38-95% buffer B gradient, a 0.1 min 95-5% buffer gradient, and a 1 min 5% buffer B wash. The resolutions of the full mass scan and tandem mass scans were 60,000 and 15,000 respectively. The full mass scan range was 500-1,200 m/z. The analysis used a 200 ms maximum injection time (IT) and 3e6 automatic gain control (AGC) for full mass scan, 100 ms maximum IT and 2e5 AGC for tandem mass scan, 0.7 m/z isolation window, a 12 loop count setting, a 1.8 kV spray voltage and a 275°C capillary temperature. Six peptides (Table 1) labeled with stable isotope arginine (13C615N4) were purchased from New England Peptides (Gardner, MA), and dissolved with 2% acetonitrile, 0.1% trifluoracetic acid at a concentration of 50 µg/mL each to generate an internal standard peptide mixture (IS). A 1 µL aliquot of this IS solution was analyzed by LC-PRM-MS to obtain the retention times and tandem MS spectra for each peptide.

#### Data analysis

After importing the raw PRM-MS data file of the IS sample into FreeStyle (version 1.5, Thermo Scientific), the extracted ion chromatographs and isotopic distributions of six targeted precursor ions were exported to .csv files, and the graphs were illustrated using GraphPad Prism (version 8.4.1, GraphPad Software). The PRM-MS result of IS solution was searched against a customized protein sequence database containing eight protein sequences downloaded in April 2020 from Uniprot database (Uniprot accessions: P9WNJ3, P9WNI7, U5WP40, H8IMX9, B5TV82, P9WNK5, B5TV81 and X8B9F3) using Thermo Proteome Discoverer (version 2.4.0.305, Thermo Scientific). The search result and raw data file were imported into Skyline software (version 20.1.0.76, MacCoss Lab, University of Washington). A spectral library was built, and the annotated tandem MS spectra were exported. The dotp and peak area ratio (endogenous version/stable isotope labeled heavy version) of each peptide peak in MGIT samples were calculated by Skyline if applicable. A peak is defined as true identification of a peptide target if its dotp is not less than 0.9, and its retention time shift compared with the pre-defined retention time of its isotope labeled version (**Table 1**) is within 1 min.

## Results

Trypsin digested CFP samples generated from four mycobacteria responsible for the majority of human mycobacterial respiratory infections (*Mtb, M. avium, M. intracellulare* and *M. abscessus*) were analyzed by LC-MS/MS to identify species-specific peptide biomarkers. Biomarker selection employed three criteria: relative abundance of the target protein in the CFP samples, a high yield of tryptic peptide production during digestion, and the presence of protein homologs in *Mtb, M. avium, M. intracellulare, M. abscessus* and *M. marinum*.

LC-MS/MS analysis identified 741, 144, 351, 50 and 236 mycobacterial proteins, respectively, in digests of *Mtb, M. avium, M. intracellulare, M. kansasii* and *M. abscessus* CFP samples (**Supplementary Table 1**), where the large number of Mtb protein identified may reflect the difference in their culture and concentration methods. *M. kansasii* CFP had the fewest proteins identified during this analysis, and the ranking protein identification scores produced by the Sequest HT searching engine included five ESAT-6-like proteins and one PE domain-containing protein associated with the ESX-1 and ESX-5 secretion systems. Substrates of the ESX-1 and ESX-5 section systems were also detected in the *Mtb* CFP digest, and substrates of ESX-5 were identified in CFP digests from *M. avium* and *M. intracellulare*, which lack ESX-1 secretion. Neither ESX-1 nor ESX-5 proteins were found in CFP digests of *M. abscessu*s, which does express either pathway. To cover the most prevalent species MAC and *M. kansasii*, and to achieve discrimination of them with *Mtb*, M. abscessus was not in the final list of this method, because it shows poor conservation in its genome when comparing it to other prevalent species. Another protein/peptide marker is needed to identify *M. abscessus*. However, the MGIT samples from *M. abscessus* infected patients were involved in this study to evaluate the method specificity.

ESX-1 substrates ESAT-6/CFP-10 and ESX-5 substrates EsxM and EsxN were detected in the CFP digests of several of these species. Since CFP-10 and ESAT-6 form a heterodimer during secretion, as do EsxM and EsxN, these proteins should exhibit a 1:1 ratio in CFP samples isolated from species that express the ESX-1 and ESX-5 secretion systems, respectively. Both dimer components should thus be useful as MS targets for species identification; however, since only CFP-10 and EsxN were consistently detected in our samples, we focused our analysis on these two targets to distinguish *Mtb*, MAC and *M. kansasii*.

We employed the BLAST algorithm to search the NCBI non-redundant protein sequence database to identify protein homologs of *Mtb* CFP-10 and EsxN among different NTM species. CFP-10 and EsxN homologs and paralogs were detected in multiple mycobacterial species, including MAC, *M. kansasii*, as well as less prevalent human pathogens such as *M. marinum, M. gordonae, M. szulgai, M. chimaera, M. ulcerans, M. xenopi, M. scrofulaceum* and *M. mucogenicum* (**Supplementary Table 2**). The ESX-1 substrate CFP-10/esxB is typically encoded by a single copy gene, and *Mtb* and *M. kansasii* have a single CFP-10 gene, but *M. marinum* expresses two CFP-10/*esxB* genes that encode slightly different protein sequences, and MAC species do not express CFP-10. However, the ESX-5 substrate EsxN has four paralogs, since a four-gene region of the ESX-5 system that contains EsxN and EsxM and a flanking *pe/ppe* gene pair has undergone four duplication events in the *Mtb* genome at sites distant from the ESX-5 locus (16, 17). Variable number duplication events for this loci are also present in the genomes of *M. kansasii, M. avium* and *M. intracellulare*. Since potential sequence variation among these EsxN paralogs could confound the ability of an EsxN-derived peptide to serve as a species-specific biomarker, we analyzed the sequence variation among EsxN and its paralogs in *Mtb* and our target NTM species by blasting the gene sequence of *Mtb* EsxN against the reference genomes of each NTM species. All chromosomal hits’ translated protein sequences were retrieved from NCBI and aligned using Jalview software to identify amino acid variations that could discriminate multiple species.

**Table 2.**
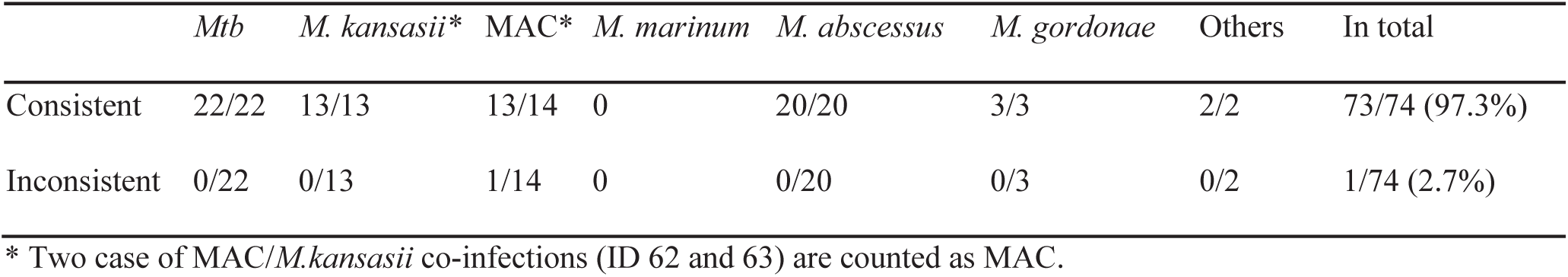
Evaluation of the PRM-MS method by comparing it with established clinical tests.

*Mtb* EsxN paralogs demonstrated 92.6 – 97.9% amino acid identity with EsxN, and differed at eight positions, including one tryptic cleavage site; however, EsxN homologs and paralogs in target NTM species exhibited greater sequence conservation. Three *M. kansasii* genes homologous to *Mtb* EsxN (MKAN_RS07975, MKAN_RS00200 and MKAN_RS04510) coded for identical protein sequences that matched 8 UniProt KB protein database entries. When searching the genome of *M. avium* EsxN genes, *Mycobacterium avium subsp. avium* ATCC 25291 that we applied for CFP digestion and protein identification by LC-MS/MS has its whole genome sequenced, but no sequence data is available in GenBank. We therefore searched its closest genome *Mycobacterium avium* strain HJW. Two EsxN paralogs were found in this genome with locus tags DBO90_RS07495 and DBO90_RS15545. They share the same protein sequence. A similar situation was observed in the genome of *M. intracellulare* ATCC 13950, which is reported to contain two genes OCU_RS30235 and OCU_RS38290 with identical sequence. Considering that *M. avium* contains several subspecies and *M. intracellulare* contains several strains which also cause lung disease, we blasted the M. avium strain HJW EsxN paralog gene DBO90_RS07495 against *Mycobacterium avium* complex (MAC) (taxid:120793) using NCBI nucleotide blast tool. We focus on *M. intracellalure* and *M. avium* in MAC because they are the major species causing human lung disease, and there is no prognostic or treatment advantage for the routine separation of MAC isolates into *M. avium* or *M.intracellulare* (18). The result showed that 54 chromosomal gene hits had at least 93.68% identity with the query sequence, and twelve plasmid hits had 88.18-89.9% identity with the query sequence (**Supplementary File 1**). Most *M. avium* subspecies have two hits to the query sequence, indicating two copies of EsxN genes. However, *Mycobacterium avium subsp. hominissuis* strain OCU901s_S2_2s, *Mycobacterium intracellulare* strain FLAC0181 have three hits. *Mycobacterium intracellulare subsp. yongonense* 05-1390 has four hits. When blasting the *M. marinum* strain E11 EsxN gene MMARE11_RS17590 against *Mycobacterium marinum* (taxid: 1781), five strains’ genomes were retrieved, and each genome contains four copies of this gene (**Supplementary File 2**). Based on this analysis, EsxN paralogs in *M. intracellulare* and *M. avium* type stains have identical sequence, which is also the case in *M. kansasii* and *M. marinum*. Neither CFP-10 nor EsxN gene can be found in *M. abscessus* genomes. These findings indicate that there is complete amino acid conservation among EsxN and its paralogs in *M. avium subsp. avium ATCC 25291, M. intracellulare*, and *M. kansasii*, and each NTM species should produce single mass targets for each peptide produced upon tryptic digestion of these related proteins. Sequence alignment of *Mtb* EsxN and EsxN paralogs, however, identified multiple sequence variations, including one that altered a trypsin cleavage site (**Supplementary Figure 1**).

Five of these amino acid variations were shared by EsxI, EsxL, and EsxV, one was shared by all but EsxN, and the remaining two were shared by EsxO and EsxL and by EsxI and EsxV. Both *Mtb* EsxI and EsxV revealed an amino acid substitution that removed the trypsin cleavage site that defines the C-terminus of the EsxN-p2 peptide (**Supplementary Figure 2**) to produce a major shift in the corresponding mass targets from these paralogs. However, none of these variations introduced sequence overlap with any of the aligned NTM EsxN proteins.

We then analyzed the CFP-10 and EsxN homologs and paralogs protein sequences in *Mtb, M. avium, M. intracellulare* and *M. marinum* to identify suitable peptide targets for species identification. CFP-10/EsxB is an important mycobacterial virulence factor that secreted by the ESX-1 pathway, which is expressed in *Mtb, M. kansasii* and *M. marinum*, but not *M. avium* or *M. intracellulare*. Two of five CFP-10 tryptic peptides (CFP10-p2 and -p4) could differentiate CFP-10 produced by *Mtb* and *M. kansasii*, and both peptides were detected in the *Mtb* CFP-10 digest, suggesting that both are suitable for species discrimination (**Figure 1A**). We selected CFP10-p2 for antibody generation due to its greater peak area in the LC-MS/MS analysis of the *Mtb* CFP-10 digest (**Figure 1C**).

**Figure 1.**
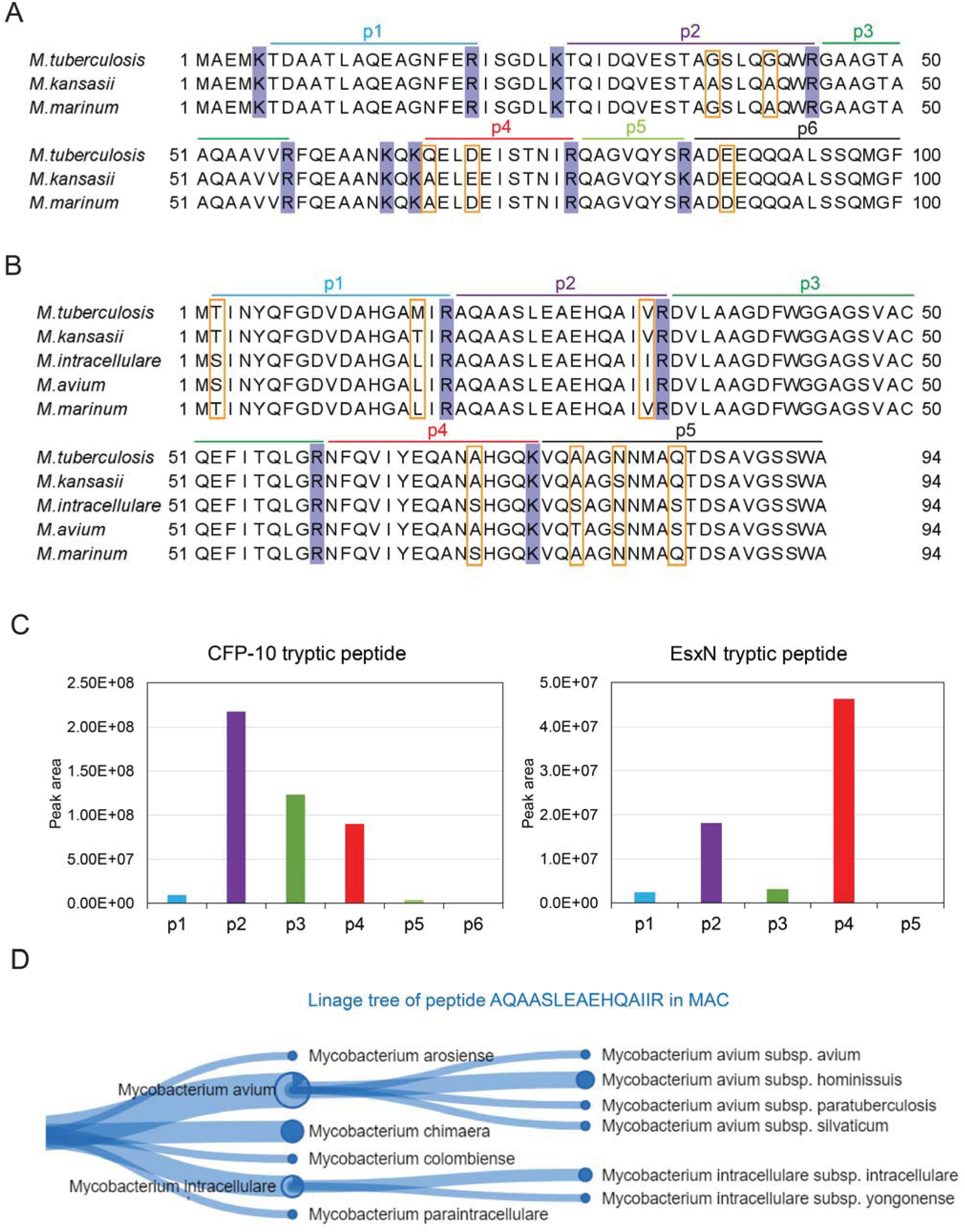
Sequence alignment of EsxB and EsxN among five mycobacterial species and mass target selection. three EsxB protein sequences (A) and five EsxN paralog protein sequences (B) were retrieved from the Uniprot database. Arginine (R) and lysine (K) are shadowed in blue, and all varied amino acids are highlighted with orange boxes. Tryptic peptides are indicated with colored lines and denoted as p1-p6. (C) The relative abundances of tryptic peptides of EsxN and CFP-10 based on the LC-MS/SM analysis of Mtb culture filtrates. The peptides are labeled as p1 to p6 according to the assignment in (A) and (B). Peak area of each peptide is shown. (D) Linage tree of EsxN peptide from MAC showing its species origin.

*Mtb* EsxN contains four tryptic cleavage sites that are conserved in EsxN homologs from EsxN and its paralogs in *M. avium, M. intracellulare*, and *M. kansasii*. EsxN-p1 could differentiate *Mtb* EsxN from homologs produced by the other three slow-growing NTM species, EsxN-p2 could differentiate *Mtb* and *M. kansasii* EsxN from holologs produced by *M. avium* and *M. intracellulare* (**Figure 1B**). EsxN-p3 was identical among all four species. EsxN-p4 could differentiate the two MAC species but could not discriminate MAC from the other analyzed species, and EsxN-p5 could distinguish all four species but was not detected in the *Mtb* CFP digest (**Figure 1C**). Based on this analysis, we excluded EsxN-p3 and p5 from further consideration as species-specific peptide biomarkers. EsxN-p2 shows the highest specificity to MAC EsxN. *Mtb* has multiple reported sub-lineages and strains. To further validate these mass targets in *Mtb*, we blasted the Mtb H37Rv CFP-10 gene sequence against *mycobacterium tuberculosis* (taxid:1773). One hundred hits were obtained, 97 were from different *Mtb* strains and 3 were from *Mtb* variant bovis (**Supplementary File 3**). Single region was matched in each genome, indicating single copy of CFP-10 gene, and 100% identity of CFP-10 sequences were observed among all hits. However, the same blast analysis of *Mtb* EsxN gene showed five or six regions matches in the 100 genomes, indicating 5-6 cpoies of EsxN genes, which is expected to be the EsxN paralogs (**Supplementary File 4**). To confirm the existence of EsxN mass targets among these *Mtb* strains, we searched the protein coding sequence of another two *Mtb* virulent clinical isolate, i.e., CDC1551 and HN878 for the EsxN paralogs. It showed that strain CDC1551 contained five EsxM paralogs in its genome with adjacent EsxN paralogs, while strain HN878 contained four EsxM and EsxN paralogs. Theoretically, when focusing on fully tryptic peptides with M+H between 800 and 3,000, three mass targets from EsxN paralogs, i.e, 1605.87, 1593.83 and 1577.84 can be observed in strain CDC1551, while two mass targets from EsxN paralogs, i.e., 1605.87 and 1619.89 can be observed in strain HN878 (**Supplementary Table 3**). The EsxN paralogs EsxO was also detected in the *Mtb* culture filtrate digest. However, the two amino acid difference between this region of EsxN and EsxO would produce a 12 Dalton mass shift that would produce two separate mass targets (1593.83 and 1605.87). Therefore, the mass targets 1593.83, 1605.87 and 2004.14 are selected to cover all three *Mtb* strains.

To further validate the existence of EsxN-p2 in multiple MAC subspecies/strains, we conducted tryptic peptide analysis of MAC EsxN-p1, p2 and p4 using the Unipept webtool (19). The results indicated that they could cover most prevalent MAC species and subspecies (**Figure 1D** and **Supplementary Figure 1**). However, the EsxN-p1 and EsxN-p4 of *M. avium* and *M. intracellulare* are shared by multiple species including MAC, the prevalent *Mtb* complex species and *M. kansasii*. Interestingly, one *Mtb* complex species *M. yongonense* TKK-01-0059 (formally known as *Mycobacterium sp.* TKK-01-0059) has its EsxN paralogs sharing EsxN-p2 with MAC. Whole-genome sequence comparison procedures showed that *M. yongonense* should be considered a member of the species *M. intracellulare* (20). EsxN-p2 is thus the most specific peptide marker of MAC.

The existence of EsxN paralogs makes it possible to observe amino acid variations in EsxN-p2 in distinct MAC subspecies and strains. We therefore retrieved the protein sequences of all chromosomal hits after blasting *M. avium* EsxN gene against MAC genomes and aligned them to identify amino acid variations in the EsxN-p2 region. *Mycobacterium intracellulare* strain FLAC0181 has two EsxN-p2 sequences, AQAASLEAEHQAIIR and TQAATLEAEHQAIIR. *Mycobacterium intracellulare subsp. yongonense* 05-1390, and *Mycobacterium avium subsp. hominissuis* JP-H-1 have three different sequences in the targeted region, including AQAASLEAEHQAIVR, AQAASLEAEHQAIIR and TQAATLEAEHQAIIR. *Mycobacterium avium subsp. hominissuis* strain MAH11 has one sequence AQAASLEAEHQAIIR, whereas strain MAC109, OCU901s_S2_2s and H87 have both AQAASLEAEHQAIVR and AQAASLEAEHQAIIR. *Mycobacterium avium subsp. paratuberculosis* strain MAPK_JJ1/13, MAPK_JB16/15, MAP/TANUVAS/TN/India/2008, MAPK_CN4/13 and E1 have two copies of EsxN genes, one has the sequence AQAASLEAEHQAIIR, the other has a mutation of tryptic site on the N-terminal of this region (R/C), resulting in the loss of this MS identifiable mass target. Other strains in *Mycobacterium avium subsp. paratuberculosis* such as strain E93, MAP4 have single sequence AQAASLEAEHQAIIR. Therefore, the sequence region that codes the MAC EsxN-p2 is highly conserved among MAC subspecies and strains. In total, four different sequences with the same number of amino acids as our peptide target in the targeted regions are identified, including AQAASLEAEHQAIVR, AQAASLEAEHQAIIR, TQAATLEAEHQAIIR and AQAASLEAEHQTIIR. The sequence AQAASLEAEHQAIIR is shared by all MAC species. In *M. marinum*, all EsxN paralogs shares their EsxN-p2 sequence as AQAASLEAEHQAIVR. Therefore, EsxN-p2 could not discriminate *Mtb* with *M. kansasii* and *M. marinum*.

Based on this analysis, we selected six peptides as PRM-MS targets (**Table 1**) that represented three species-selective peptides from CFP-10-p2 and EsxN-p2 and its paralog equivalents that could distinguish *Mtb*, MAC, *M. kansasii* and *M. marinum* in CFP digests when analyzed in combination, and generated peptide specific antibodies to the *Mtb* variants of the CFP-10-p2 and EsxN-p2 peptides.

These antibodies were then employed to enrich their EsxN-p2 and CFP-10-p2 peptides from MGIT cultures inoculated with clinical samples time to positivity ranges of 2 days to 32 days, where species identification was determined by established methods (**Figure 2**).

**Figure 2.**
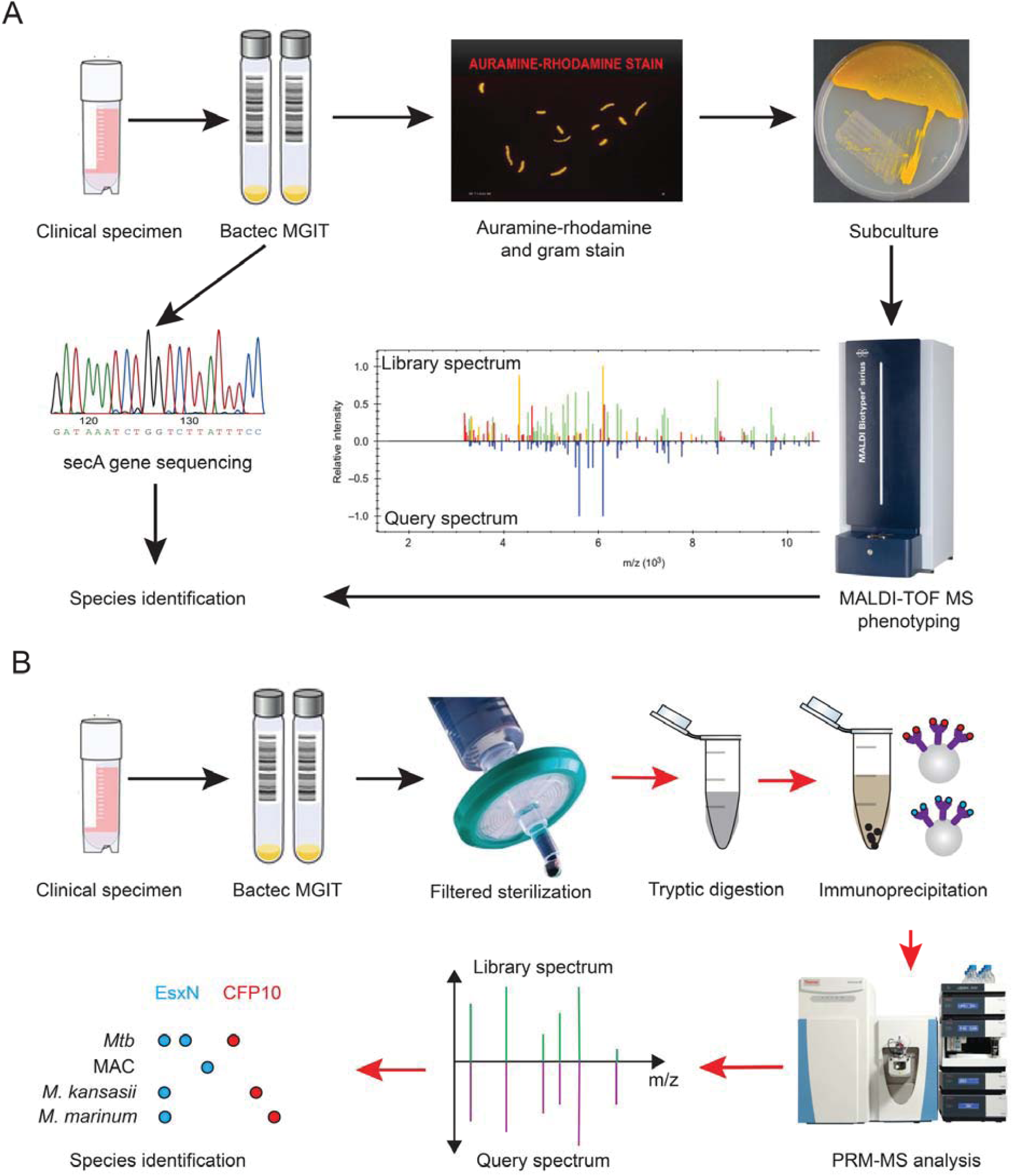
Schematic of the routine workflow of mycobacteria species identification in clinical labs (A) and species identification through the proposed method (B). Red arrows in (B) show the procedure that can be conducted at a BSL-2 lab. Tandem MS spectra of peptide targets are annotated based on their identifications after database search and used to build a spectral library to identify corresponding peptides in MGIT samples.

The PRM-MS method was established by analyzing the LC-MS/MS profiles of mixed samples of synthetic CFP10-p2 and EsxN-p2 peptides generated with a carboxy-terminal stable-isotope arginine label. All three EsxN-p2 targets were separated by LC (**Supplementary Figure 3**), although two peaks were observed for the Mtb EsxL-p2 peptide, with the second peak deriving from the third isotopic peak of the EsxO-p2 IS peptide (**Table 1**), and the most intense peaks in its MS/MS spectrum matched this peptide (**Supplementary Figure 4**). However, despite this overlap both peptides were unambiguously identified by PRM-MS (**Figure 3**) by setting a high threshold for the peptide similarity score (dot product [dotp]: 0.9) in the Skyline analysis software. PRM-MS identified single EsxN and EsxO peaks meeting this threshold in a clinical MGIT samples from a patient with an active *Mtb* infection (**Figure 4A-B**), and similar results were found upon analysis of MGIT samples from patients with MAC and *M. kansasii* infections (**Figure 4C-D**). Similar LC results were also found for the analyzed CFP-10 mass targets (**Supplementary Figure 3**). Additional peaks detected for EsxN-p2, or its paralog peptides, in some samples resulted from mass similarity, despite the narrow window (0.7 m/z) used for precursor ion isolation. Reducing the isolation window width could resolve this issue, but might also decrease sensitivity. However, our current results indicated that the dotp value for the candidate mass targets may be sufficient to determine the correct mass target peak.

**Figure 3.**
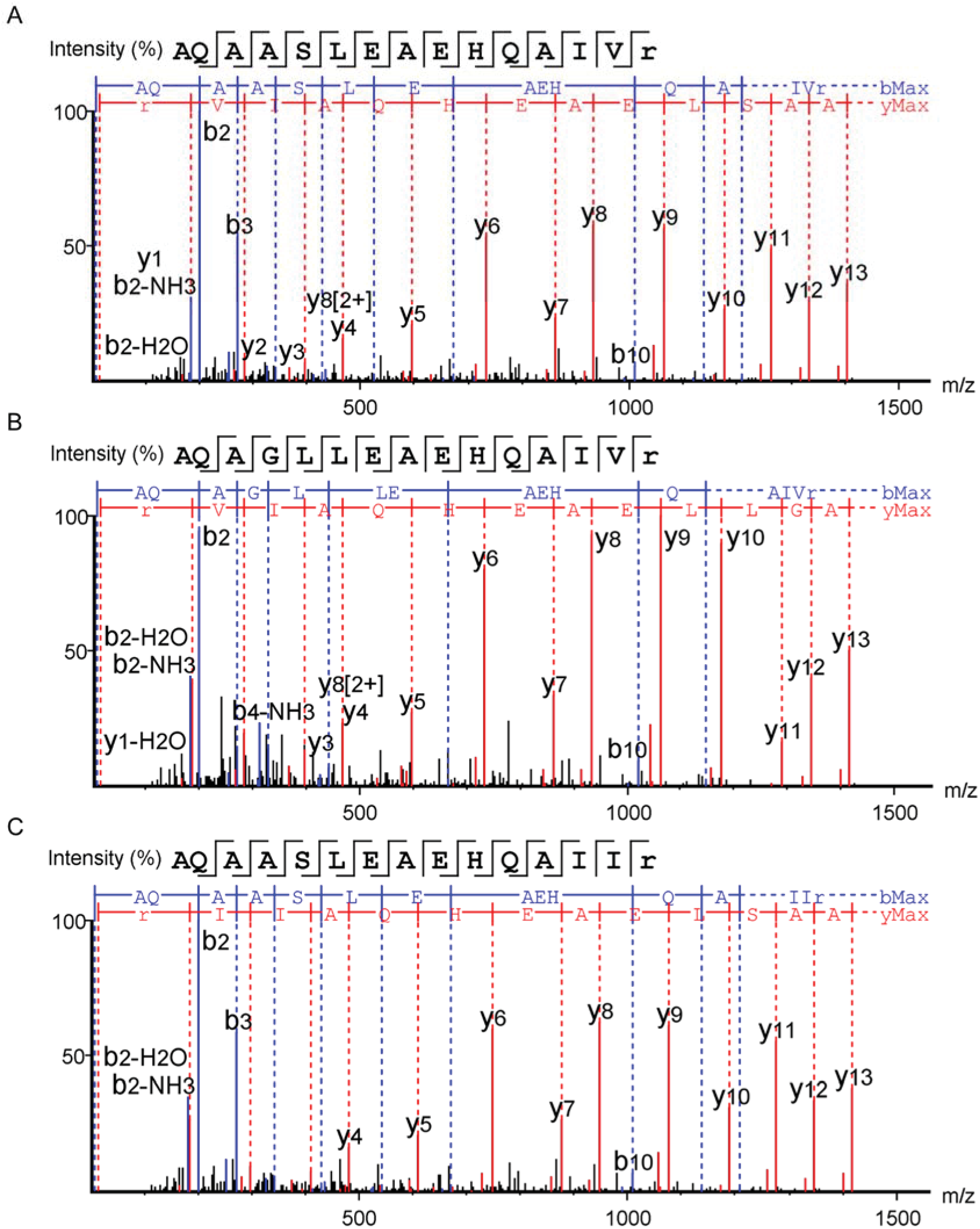
Tandem MS spectra with fragment annotations of the three peptide targets from EsxN paralogs. (A) *Mtb* and *M. kansasii* EsxN peptide AQAASLEAEHQAIVR. (B) Mtb EsxO peptide AQAGLLEAEHQAIVR. (C) MAC EsxN peptide AQAASLEAEHQAIIR.

**Figure 4.**
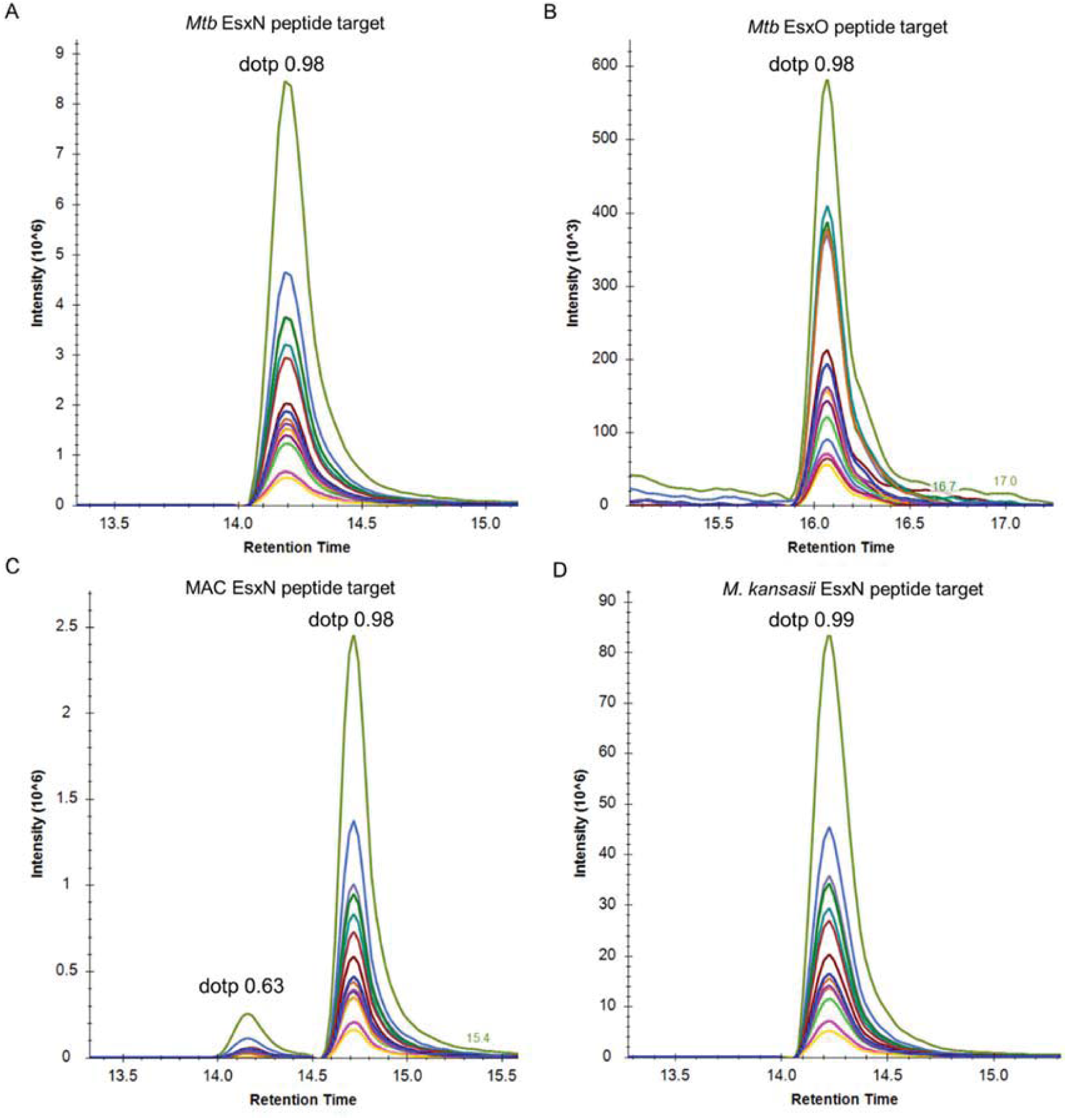
Extracted ion chromatograms of the two peptide targets from *Mtb* EsxN (A), *Mtb* EsxO (B), MAC EsxN paralogs (C) and *M. kansasii* EsxN paralogs (D). Co-elution of their fragments are shown by their ion chromatograms extracted from PRM spectra of MGIT samples. They are colored as indicated. Dotp value of the peaks at expected retention times are labeled.

PRM-MS analysis of a *Mtb-*positive MGIT culture detected EsxN-p2 and EsxO-p2 (1593.83 and 1605.87 m/z) and CFP10-p2 (2004.14 m/z) mass targets, a *M. kansasii*-positive MGIT sample detected mass targets for CFP10-p2 and EsxN-p2 and/or its identical paralogs (2032.01 and 1593.83 m/z), while a MAC-positive MGIT sample detected mass targets for and EsxN-p2 and/or its paralogs (1607.85 m/z) (**Supplementary Figures 5-7**).

To evaluate the diagnostic performance of our PRM-MS assay approach, we analyzed 82 MGIT samples from 74 patients, and found that PRM-MS had an overall consistency of 98.6% (73/74) results determined by other analysis methods. This analysis revealed robust PRM-MS diagnostic sensitivity for *M. kansasii* (12/13; 92.3%), *Mtb* (22/22; 100%), and MAC (14/14; 100%), with high specificity (25/25; 100%) (**Table 2**).

Patients in the negative control group were infected with the three rapidly growing species of mycobacteria (*M. abscessus, M. fortuitum* or *M. massiliense*), which do not produce mass targets recognized by our assay, or the slow-growing mycobacteria *M. gordonae*. Notably, all three patients with *M. gordonae* infections were found to express a CFP-10 peptide not targeted by PRM-MS, but was recognized due to its identical mass and a 0.3-min retention time shift when compared to the *M. kansasii* CFP-10 peptide (**Figure 5A-B**). This peptide had a low dotp when compared to the library spectrum of *M. kansasii* CFP-10, and a database search for the PRM spectra found this spectrum matched the *M. gordonae* CFP-10 peptide (**Figure 5C**).

**Figure 5.**
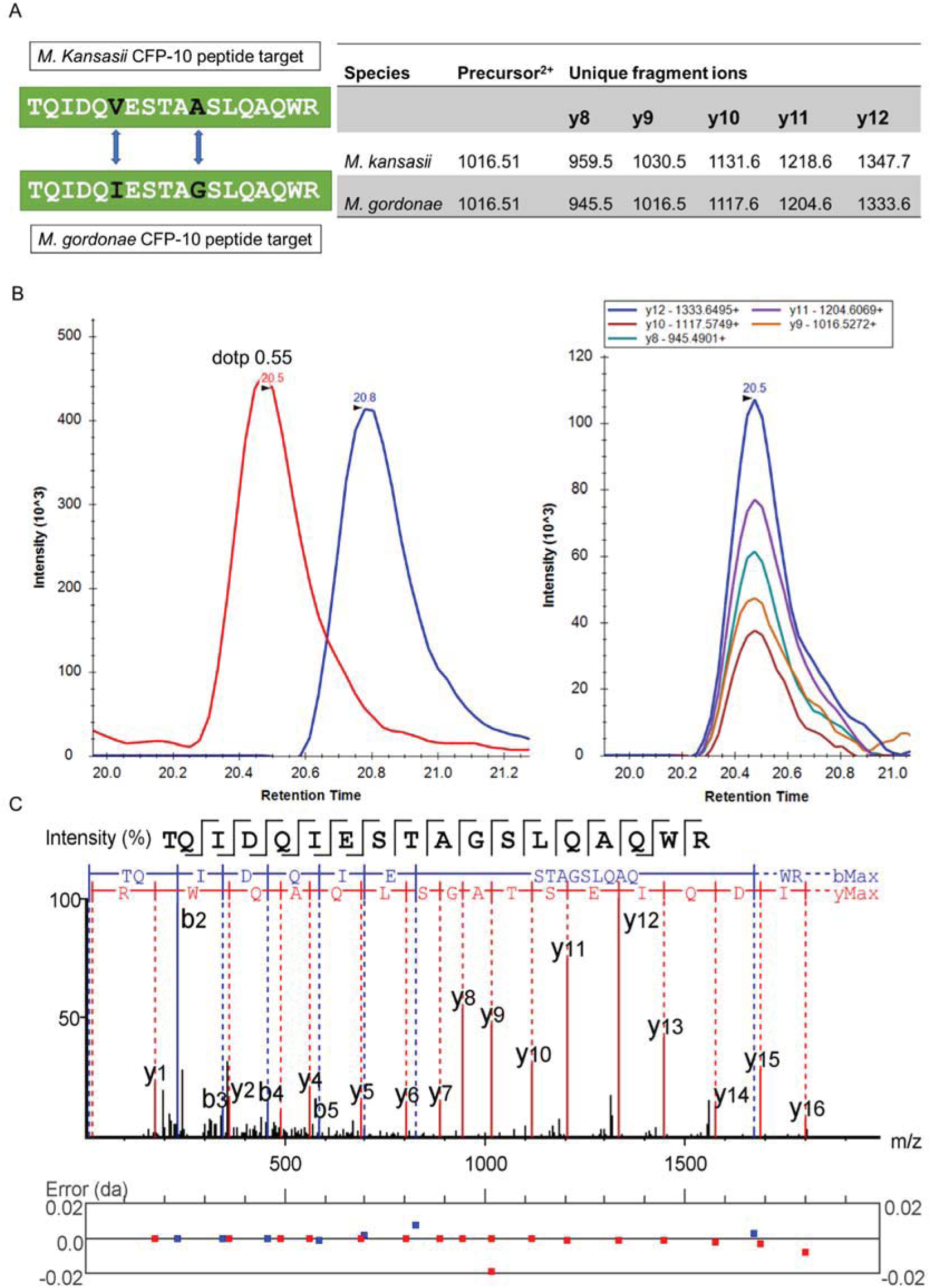
PRM-MS result of an MGIT sample from a patient with *M. gordonae* infection. (A) The sequences of CFP-10 peptide target from *M. kansasii* and *M. gordonae.* Two amino acid variations are highlighted. The precursor m/z and distinguishable fragment ions are listed in the right table. (B) Extracted ion chromatograms of *M. kansasii* CFP-10 peptide TQIDQVESTAASLQAQWR in both light (red) and heavy (blue) versions. Note there is a 0.3-minute retention time shift between the light and the heavy peptide peaks. In the right side, extracted ion chromatograms of five distinguishable fragment ions of peptide *M. gordonae* CFP-10 peptide TQIDQIESTAGSLQAQWR are shown. The chromatograms are colored as indicated and their m/z are shown in the top box. (C) Annotated tandem MS spectrum of this peptide identified as the tryptic peptide from *M. gordonae* CFP-10. All matched peaks are colored and labeled with their fragment ion names. Mass error of each matched ion is shown in the plot below.

Since *M. gordonae*-positive MGIT samples are primarily the result of environmental contamination (21), detection of this species may indicate sample contamination rather than infection.

The MAC EsxN-p2 peptide (1607.85 m/z) was detected in MGIT samples from all but one patient with a MAC infection (13/14; 92.9%) with a second target (1593.83 m/z) detected in 3 of these positive samples (**Figure 6A-B**). Two MAC species, *M. chimaera* (strain CDC 2015-22-71) and *M. intracellulare yongonense* 05-1390 contain EsxN paralogs (UniProt KB A0A1Y0SZB6 and S4ZJN8) that can generate this mass target (**Supplementary Table 3** and **Supplementary File 1**). A third mass target (2032.2 m/z) detected in samples from two of the 13 patients (**Figure 6C**), appears to be unique to *M. kansasii, M. gastri, M. innocens* and *M. attenuatum* (22), based on tryptic peptide analysis using Unipept webtool, suggesting these the two patients are have MAC and *M. kansasii* co-infections, consistent with molecular diagnosis results for these patients (**Supplementary Table 4**). As mentioned above, the 2032.2 mass target could be detected in both *M. kansasii* and *M. gordonae* infected samples. However, the *M. kansasii* CFP-10-p2 unique fragment ions were observed in these samples (**Figure 6D**), confirming the existence of *M. kansasii* instead of *M. gordonae*.

**Figure 6.**
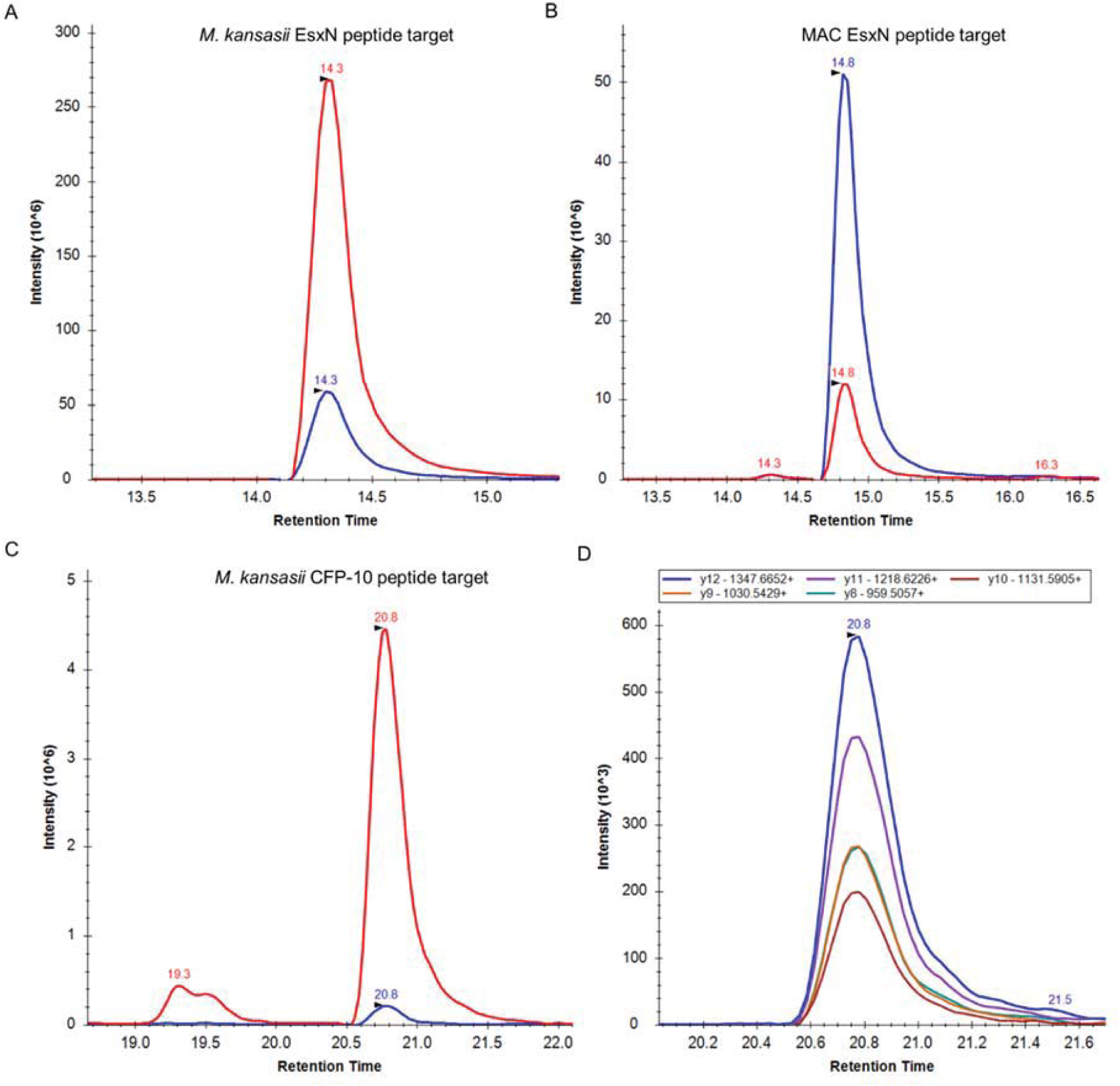
PRM-MS result of an MGIT sample from patients with *M. avium* and *M. kansasii* co-infections. The co-infections were confirmed by identifying the *M. kansasii* EsxN peptide target (A), *M. kansasii* CFP-10 peptide target (B) and MAC EsxN peptide target (C) simultaneously. The identified peptide peaks are shown with their light version (endogenous peptide) colored red and heavy version (stable isotope labeled peptide) colored blue in their extracted ion chromatograms. (D) Extracted ion chromatograms of the five fragment ions from *M. kansasii* CFP-10 that can discriminate it with *M. gordonae* CFP-10 peptide.

Finally, EsxO-p2 (1605.87 m/z) was detected in the majority (14/22; 63.6%) of MGIT cultures from patients with active *Mtb* infections. Four of the subjects without EsxO signal were resistant to at least one anti-TB drug, and another three were sensitive to more than one first-line drug (**Supplementary Table 4**), suggesting a potential relationship with drug resistance, possibly resulting from drug resistant *Mtb* strains that exhibit differential EsxO expression or mutations.

## Discussion

Currently, weakly sensitive microscopic assays and time-consuming culture methods are used to diagnose and monitor mycobacterial infections. Mycobacterial cultures, the diagnostic gold standard, requires up to 6 weeks to produce a final report. Bacterial proteins secreted in the early phase of mycobacterial growth are promising candidate markers for early, specie-specific diagnosis of mycobacterial infections. We selected the ESX-5 substrate EsxN and the ESX-1 substrate CFP-10 as our target in this work, due to their high abundance in CFP and the essential roles they play in bacterial growth and evading immune surveillance (23). We next selected an *Mtb* EsxN peptide region and an *Mtb* CFP-10 peptide region that were consistently detected in all samples with EsxN and EsxB signal and that demonstrated the ability to distinguish among several species as mass targets, and generated antibodies against these peptides were used to develop an IP-based MS assay. In this approach, the specificity provided by antibody capture of targeted tryptic peptides and their tandem MS spectra detection by PRM-MS enabled accurate species discrimination. This assay could discriminate cultures of *M. avium* and *M. intracellulare*, which are responsible for the majority of human mycobacterial respiratory infections, from those of *Mtb, M. kansasii* or *M. gordonae*, which could also be distinguished by this method. In three MGIT samples (ID 55, 66 and 69), we observed both 1593.83 and 1607.85. They are identified as *M. avium* infections by established methods. As mentioned above, *mycobacterium avium subsp. hominissuis* strain MAC109, OCU901s_S2_2s and H87 may express both targets. Therefore, the identification of mass target 1593.83 in these samples may be explained by the infection of these specific strains. It is difficult to discriminate *M. intracellulare* with *M. chimaera* using established methods (24, 25). This explained that four MGIT samples with IDs 63, 67, 74 and 75 were identified as *M. intracellulare/M. chimaera*. By using the proposed method, *M. chimaera* cannot be discriminated with *Mycobacterium intracellulare* strain FLAC0181 or *Mycobacterium intracellulare subsp. yongonense* 05-1390, because both 1593.83 and 1607.85 mass targets are expected to be expressed by these species/strains. Therefore, these samples were identified as MAC, and more peptide targets are needed to further identify the species. The mass targets of *M. marinum* were included in the current assay, however, this species mainly cause skin infection (26).The CFP-10 mass target of *M. marinum* was not observed in any MGIT sample, indicating a high species specificity of this peptide target.

The species identification accuracy of the proposed method is based on high resolution tandem mass spectra, which can even discriminate the isomers such as CFP-10 peptides from *M. gordonae* and *M. kansasii* with two amino acid variations. The tandem MS spectra of internal standard peptides are used to build a spectral library. Similar to the mycobacteria library utilized in commercial MALDI-TOF phenotyping assay, this spectral library is a reference and the similarity-based dotp value between a query spectrum and a library spectrum indicates the confidence of peptide identification. However, this peptide spectral library is more stable than the spectral library of extracted bacterial proteins used in commercial MALDI-TOF phenotyping assay, as the latter of which may be affected by the reference strain of mycobacteria utilized to build the library. The proposed method also showed the ability to identify MGIT samples with *M.kansasii* and MAC co-infections. When MGIT tubes are polymicrobial, the proposed method can achieve identification of multiple species by their distinct mass targets of EsxN paralogs and CFP-10.

In theory, it is possible to directly measure the secreted bacterial proteins in MGIT filtrates using LC-MS/MS for species identification. However, the abundant albumin existed in MGIT filtrates needs to be removed before loading the sample to LC-MS/MS. Therefore, we processed the culture filtrates with ultracentrifuge for LC-MS/MS analysis. The advantage of measuring secreted proteins without IP is that it allows identifying mycobacteria that express neither EsxN paralogs nor CFP10, such as *M. abscessus*. In this regard, it would be feasible to process an MGIT sample by both ultracentrifuge and IP to generate two fractions. An untargeted LC-MS/MS analysis of the sample processed by ultracentrifuge will achieve identification of some mycobacteria species if they express specific proteins with detectable amounts in MGIT filtrates. A targeted PRM-MS analysis of the fraction processed by IP will provide complementary result if the samples contain slowly growing mycobacteria that express EsxN paralogs or CFP-10 with a lower amount. With the increasing genome and proteome databases of mycobacteria available now, it is possible to build a protein sequence database containing multiple mycobacteria species and search the tandem MS spectra of MGIT filtrates against this database for species identification. The advantage of using this sequence database instead of MALDI-MS spectra library is to identify mycobacteria unambiguously by using its unique peptides. The high-resolution MS even allows identification of homologeous peptides with single amino acid variation, as exemplified by the two EsxN peptide targets AQAASLEAEHQAI<u>V</u>R and AQAASLEAEHQAI<u>I</u>R.

A recent study provided evidence that the Beijing lineage carries a variant of esxW, that is Thr2Ala under positive selection in natural *Mtb* populations, implicating a possible regulation mechanism through substrate selection for ESX-5 secretion under certain conditions (27). The EsxW-Thr2Ala alteration was also identified in 9 of 11 patients with MDR-TB reinfection (28). It is thus reasonable to detect the mutation of esxW gene in its protein product. By applying top-down proteomics developed in recent years, measuring such a mutation of esxW in MGIT samples is achievable. Except for gene mutations, the drug-resistant *Mtb* strains may show a selective expression of EsxN paralogs in its culture media compared with that of the drug-susceptible strains. In our results, the mycobacteria in four MGIT samples from patients infected with drug resistant *Mtb* express abundant EsxN, but the expression of EsxO is undetectable. It is established that *Mtb* requires ESX-5 for virulence, and secretion of ESX-5 substrates including the EsxN paralogs is tightly controlled to avoid elimination by host immune responses (29). A functional model is proposed for the modular organization of the ESX-5 secretion system, that is selective secretion of a subset of the ESX-5 secretome (23). However, four paralogs were listed in that model, and *esxO* gene that we observed in some MGIT samples from patients with *Mtb* infections was missing. By applying the proposed IP-MS analysis, one can easily identify the selective expression of different EsxN paralogs in *Mtb* infected samples. Additional studies are ongoing to identify drug-resistance related amino acid mutations and/or selective expression of secreted proteins of mycobacteria. Once a specific mutation or a subset of secreted bacterial proteins relating to drug resistance is identified, this type of multiplexed IP-MS assay can be easily applied to identify the drug resistance phenotype.

The proposed method shows potential application for culture-independent rapid diagnosis by using blood samples considering that ESX-5 secretion system is critical for mycobacterial persistence in host cells. The ESX-5 substrates may circulate in the patient blood and play roles in modulating host immune response. Ongoing study will validate the identified ESX-5 substrates in the circulation of patients with NTM infections, which will dramatically shorten the time for diagnosing mycobacteria infections.

## Abbreviations

BSL: biosafety level
CFP: culture filtrate protein
CID: collision induced dissociation
DST: drug susceptibility tests
ESX: Esat-6 protein secretion systems
HCD: higher-energy collisional dissociation
IP: immunoprecipitation
IP-MS: immunoprecipitation coupled with mass spectrometry
LC: liquid chromatography
LC-MS/MS: liquid chromatography coupled with tandem mass spectrometry
MAC: *M. avium* complex
MALDI-TOF MS: matrix assistant laser desorption ionization-time of flight mass spectrometry
MGIT: mycobacterial growth indicator tube
MS: mass spectrometry
*Mtb*: *Mycobacterium tuberculosis*
NTM: non-tuberculous mycobacteria

---

**Supplementary Figure 1.**
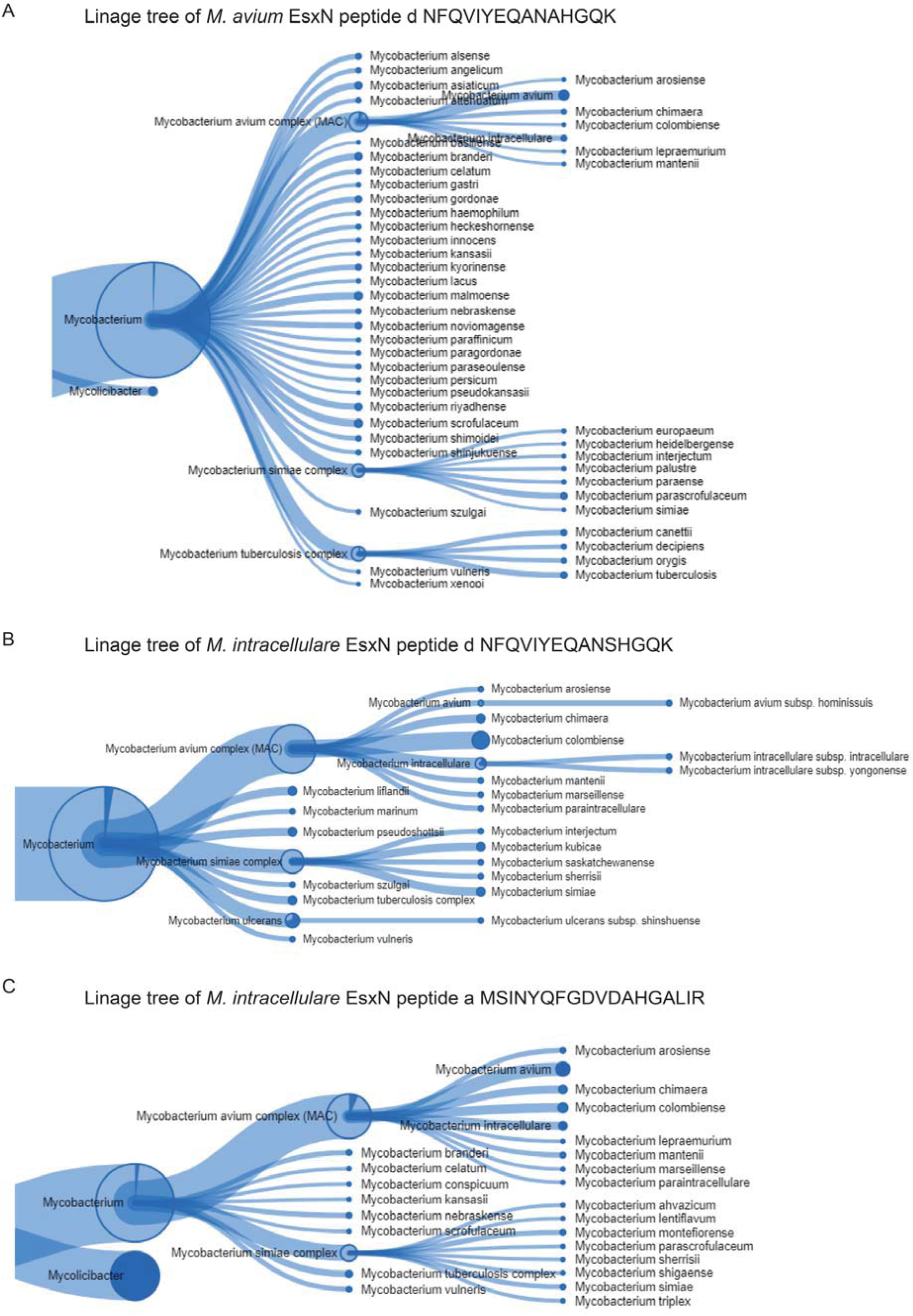
Linage tree of protein hits that contain the queried peptide sequences from MAC EsxN after analyzing them with Unipept tool.

**Supplementary Figure 2.**
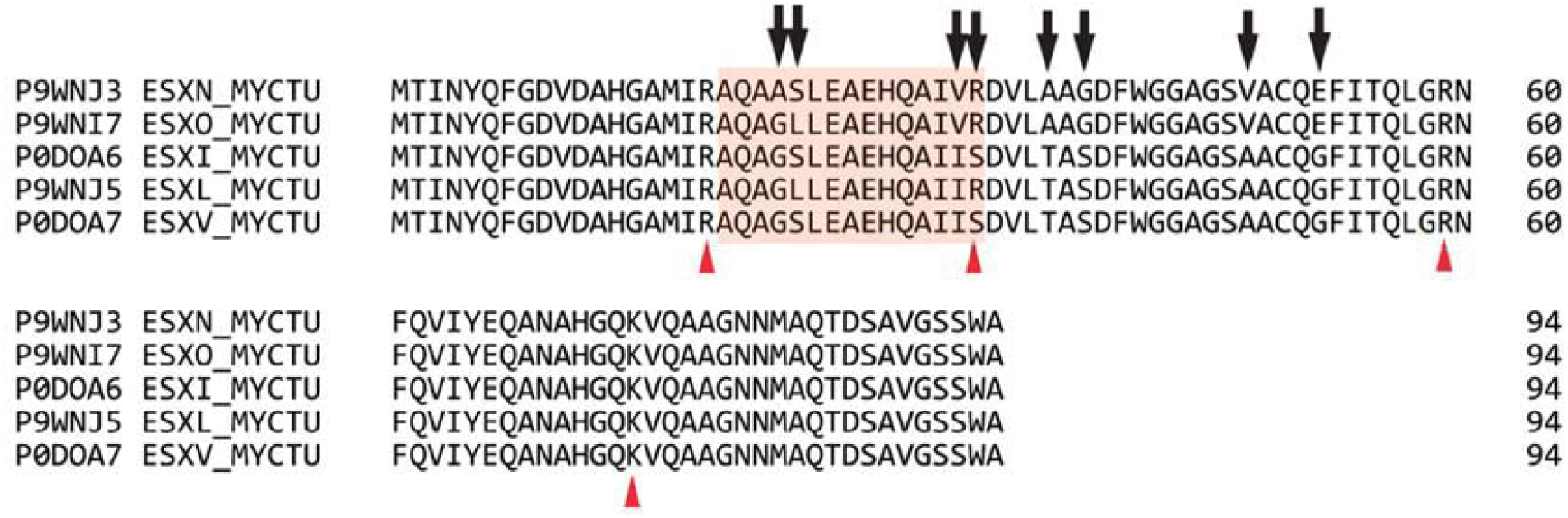
Sequence alignment of Mtb EsxN and its four paralogs. Trypsin cleavage sites are marked with arrowheads. The peptide region chosen for mass target analysis is indicated by highlighting. Aligned amino acids are marked to indicate identity (*), conservative (:) and semi-conservative (.) differences, or lack of homology (blank). Individual amino acid differences to EsxN sequence are highlighted on the corresponding peptide sequence.

**Supplementary Figure 3.**
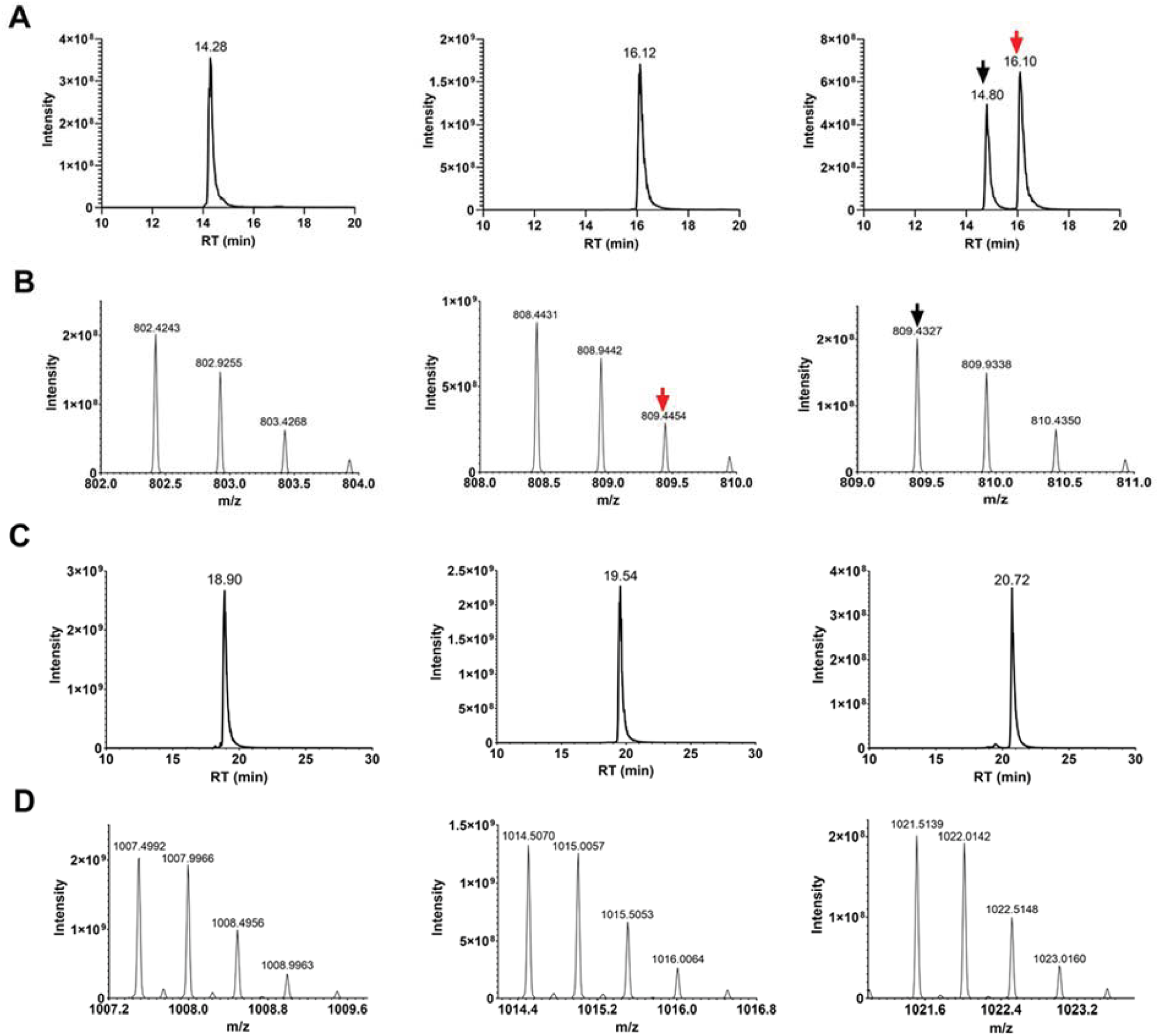
Retention times of the three peptide targets from EsxN paralos after PRM-MS analysis (A: left, AQAASLEAEHQAIVR; middle, AQAGLLEAEHQAIVR; right, AQAASLEAEHQAIIR) and from CFP10 (C: left, TQIDQVESTAGSLQGQWR; middle, TQIDQVESTAASLQAQWR; right, TQIDQVESTAGSLQAQWR). Their precursor ion isotopic distributions (B and D) are used to determine the corresponding peaks in their extracted ion chromatograms (A and C). An ambiguous peak with precursor m/z 809.4327 is observed for AQAASLEAEHQAIIR (red arrow in A), which is caused by the third isotopic peak of peptide AQAGLLEAEHQAIVR. The correct peak is indicated with black arrow.

**Supplementary Figure 4.**
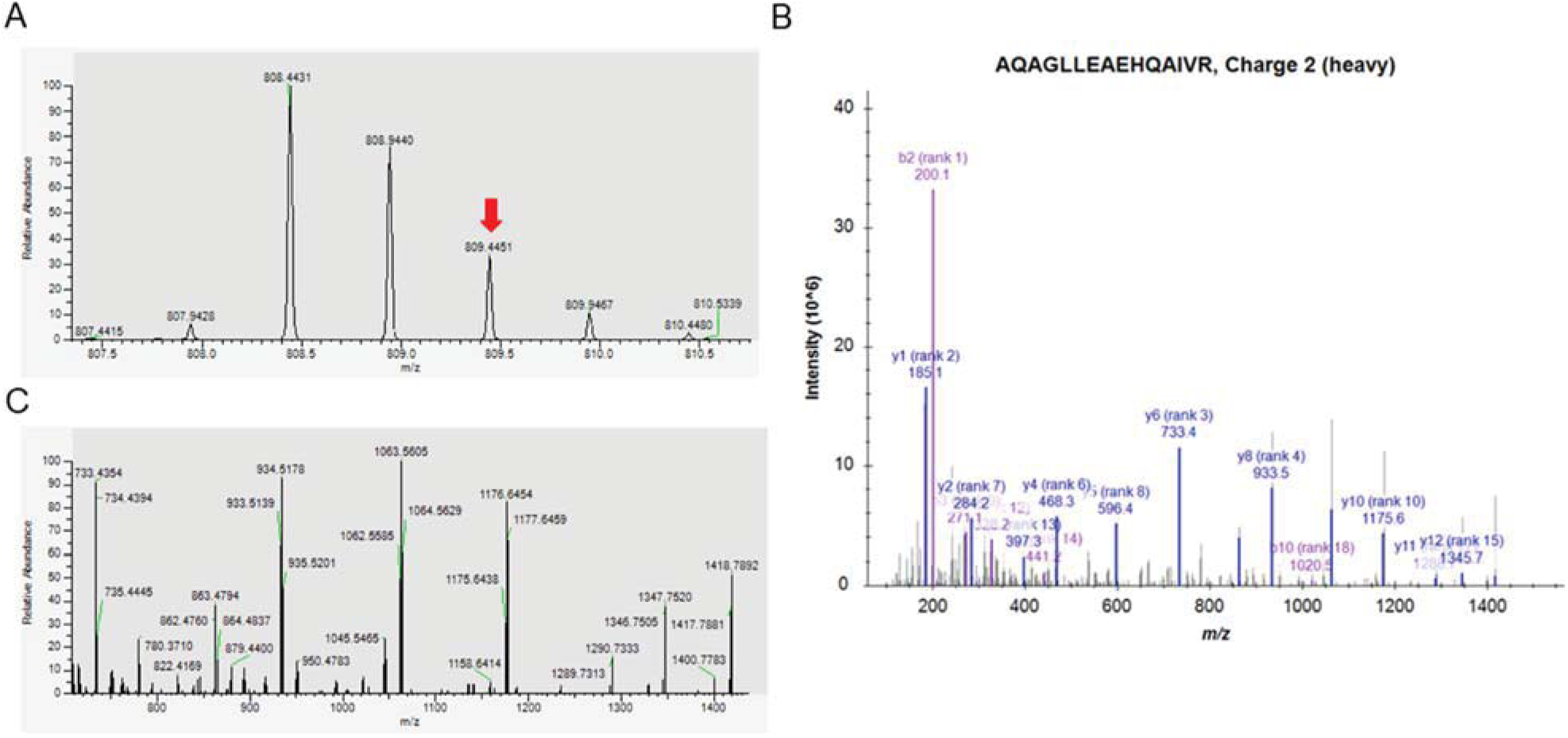
Confirmation of the ambiguous peak in Supplementary Figure 2C derived from the isotopic peak of another peptide. (A) The isotopic distribution of peptide AQAGLLEAEHQAIVR. After stable isotope labeling, its third isotopic peak indicated with red arrow is close to the precursor m/z of AQAASLEAEHQAIIR, and its tandem spectrum (B) was identified as the sequence of AQAGLLEAEHQAIVR (C). The matched fragments are labeled with their m/z and relative intensity ranks.

**Supplementary Figure 5.**
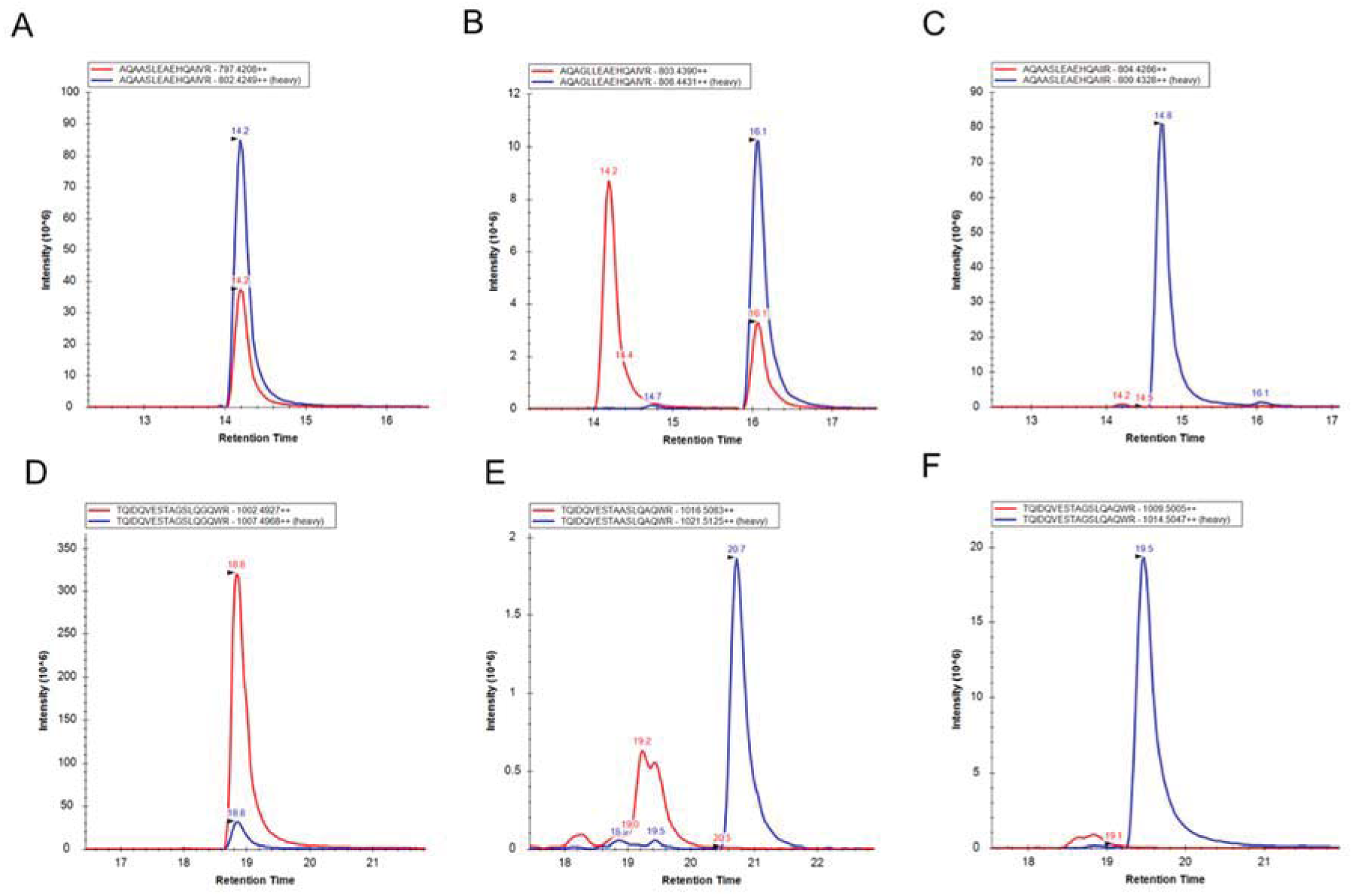
PRM-MS result of an MGIT sample from patients with *Mtb* infections. Peptide sequences are present in the top box, with their light version (endogenous peptide) colored red and heavy version (stable isotope labeled peptide) colored blue in their extracted ion chromatograms. The correct peaks are labeled with black arrowheads according to their retention times.

**Supplementary Figure 6.**
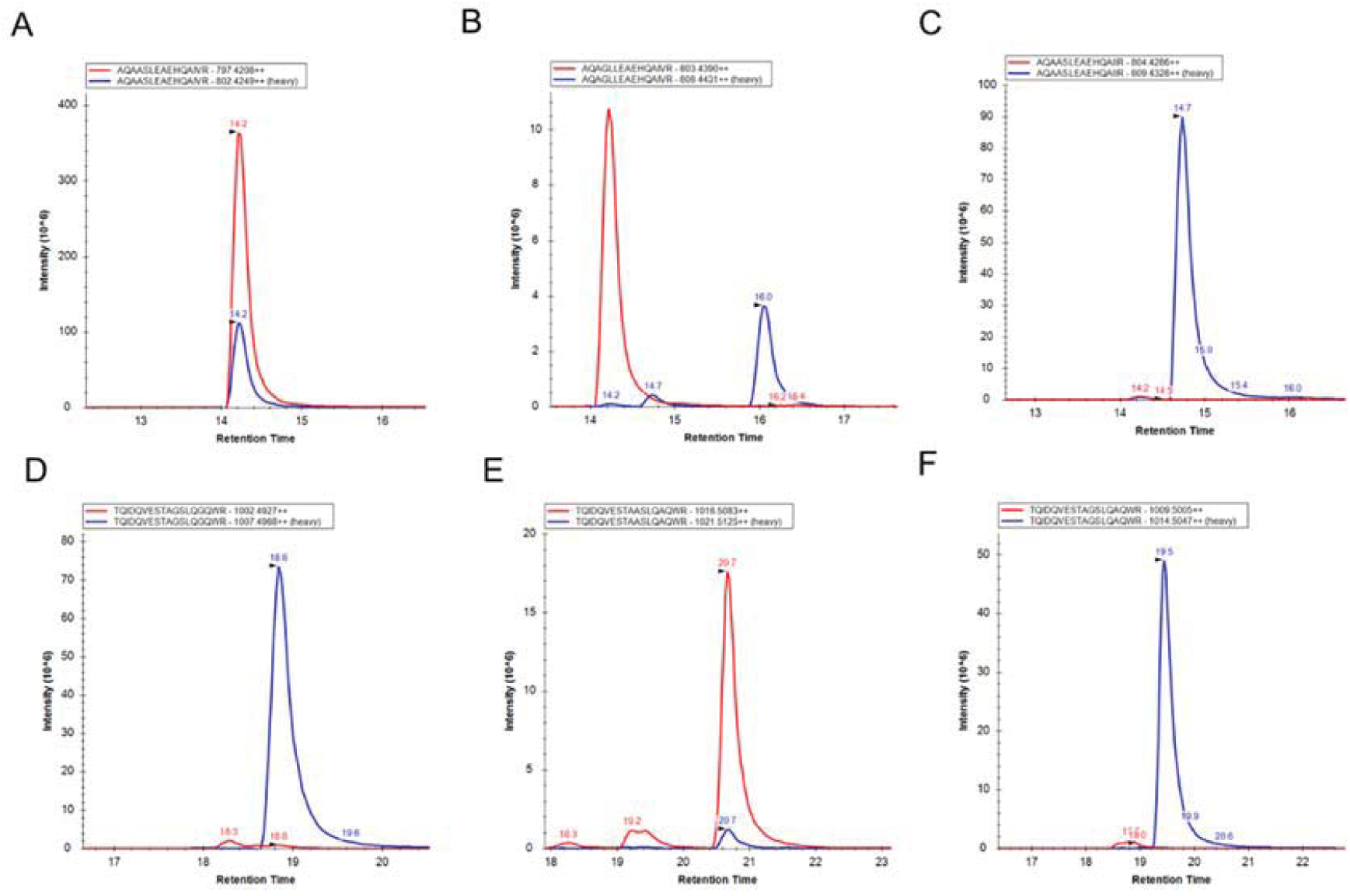
PRM-MS result of an MGIT sample from patients with *M. kansasii* infections. Peptide sequences are present in the top box, with their light version (endogenous peptide) colored red and heavy version (stable isotope labeled peptide) colored blue in their extracted ion chromatograms. The correct peaks are labeled with black arrowheads according to their retention times.

**Supplementary Figure 7.**
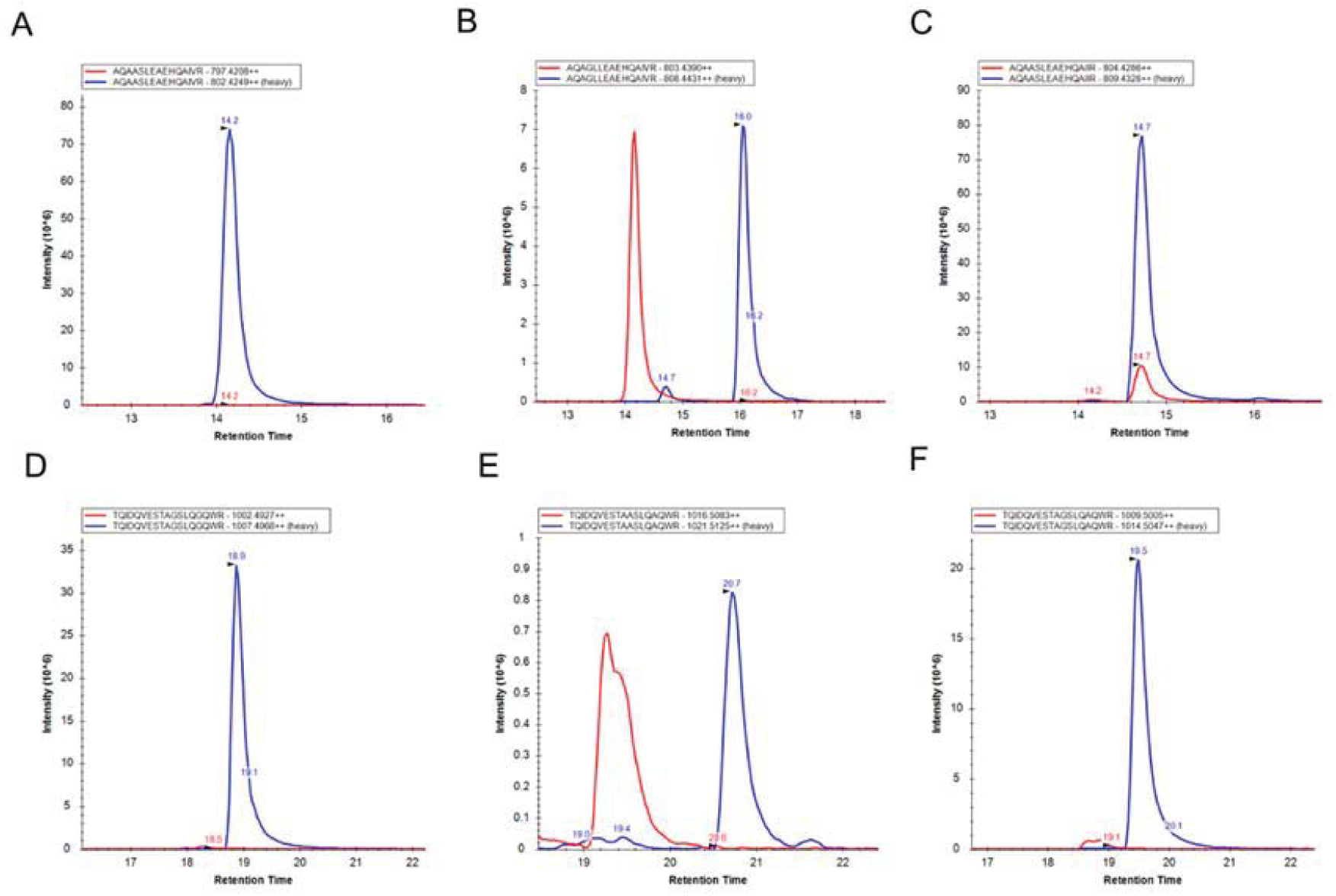
PRM-MS result of an MGIT sample from patients with *M. avium* infections. Peptide sequences are present in the top box, with their light version (endogenous peptide) colored red and heavy version (stable isotope labeled peptide) colored blue in their extracted ion chromatograms. The correct peaks are labeled with black arrowheads according to their retention times.

## Conflict of Interest

The authors declare that the research was conducted in the absence of any commercial or financial relationships that could be construed as a potential conflict of interest.

## Author Contributions

Q.S., J.F., L.Z., X.K. and Y.H. designed, performed, and analyzed the experiments. M.R., S.R. and A.Z. provided the clinical isolates MGIT samples and conducted mycobacterial identification using established methods, provided suggestions and critically read the paper. Y.H. and W.S. conceptualized and designed experiments, analyzed the results. Q.S. and C.L. wrote the paper.

## Funding

The work was primarily supported by research funding provided by NIH (R01HD090927, R01AI122932, R01AI113725 and R21AI126361-01), and Arizona Biomedical Research Commission (ABRC) young investigator award.

## Acknowledgments

The following reagents were obtained through BEI Resources, NIAID, NIH: Mycobacterium tuberculosis, Strain H37Rv and CDC1551, Culture Filtrate Proteins, NR-14825 and NR-14826.

## Data Availability Statement

The mass spectrometry proteomics data have been deposited to the ProteomeXchange Consortium via the PRIDE partner repository with the dataset identifiers PXD019069 and PXD019068 (30).

